# Metabolic reprogramming and flux to cell envelope precursors in a pentose phosphate pathway mutant increases MRSA resistance to β-lactam antibiotics

**DOI:** 10.1101/2023.03.03.530734

**Authors:** Merve S. Zeden, Laura A. Gallagher, Emilio Bueno, Aaron C. Nolan, Jongsam Ahn, Dhananjay Shinde, Fareha Razvi, Margaret Sladek, Órla Burke, Eoghan O’Neill, Paul D. Fey, Felipe Cava, Vinai C. Thomas, James P. O’Gara

## Abstract

Central metabolic pathways controls virulence and antibiotic resistance, and constitute potential targets for antibacterial drugs. In *Staphylococcus aureus* the role of the pentose phosphate pathway (PPP) remains largely unexplored. Mutation of the 6-phosphogluconolactonase gene *pgl,* which encodes the only non-essential enzyme in the oxidative phase of the PPP, significantly increased MRSA resistance to β-lactam antibiotics, particularly in chemically defined media with glucose, and reduced oxacillin (OX)-induced lysis. Expression of the methicillin-resistance penicillin binding protein 2a and peptidoglycan architecture were unaffected. Carbon tracing and metabolomics revealed extensive metabolic reprogramming in the *pgl* mutant including increased flux to glycolysis, the TCA cycle, and several cell envelope precursors, which was consistent with increased β-lactam resistance. Morphologically, *pgl* mutant cells were smaller than wild-type with a thicker cell wall and ruffled surface when grown in OX. Further evidence of the pleiotropic effect of the *pgl* mutation was reduced resistance to Congo Red, sulfamethoxazole and oxidative stress, and increased resistance to targocil, fosfomycin and vancomycin. Reduced binding of wheat germ agglutinin (WGA) to *pgl* was indicative of lower wall teichoic acid/lipoteichoic acid levels or altered teichoic acid structures. Mutations in the *vraFG* or *graRS* loci reversed the increased OX resistance phenotype and restored WGA binding to wild-type levels. VraFG/GraRS was previously implicated in susceptibility to cationic antimicrobial peptides and vancomycin, and these data reveal a broader role for this multienzyme membrane complex in the export of cell envelope precursors or modifying subunits required for resistance to diverse antimicrobial agents. Altogether our study highlights important roles for the PPP and VraFG/GraRS in β-lactam resistance, which will support efforts to identify new drug targets and reintroduce β-lactams in combination with adjuvants or other antibiotics for infections caused by MRSA and other β-lactam resistant pathogens.

**Author summary:** High-level resistance to penicillin-type (β-lactam) antibiotics significantly limits the therapeutic options for patients with MRSA infections necessitating the use of newer agents, for which reduced susceptibility has already been described. Here we report for the first time that the central metabolism pentose phosphate pathway controls MRSA resistance to penicillin-type antibiotics. We comprehensively demonstrated that mutation of the PPP gene *pgl* perturbed metabolism in MRSA leading to increased flux to cell envelope precursors to drive increased antibiotic resistance. Moreover, increased resistance was dependent on the VraRG/GraRS multienzyme membrane complex previously implicated in resistance to antimicrobial peptides and vancomycin. Our data thus provide new insights on MRSA mechanisms of β-lactam resistance, which will support efforts to expand the treatment options for infections caused by this and other antimicrobial resistant pathogens.

## Introduction

The World Health Organization (WHO) recently reported a dramatic increase in antimicrobial resistance (AMR) among human pathogens (1, 2). Exacerbation of the AMR crisis is driven by the misuse and overuse of last-resort antibiotics, the decline in new antimicrobial drugs being approved for clinical use and a lack of mechanistic understanding of AMR in bacterial pathogens (1, 2). *Staphylococcus aureus*, which is among the most challenging AMR human pathogens, can cause a variety of infections. Skin and soft tissue infections can be localised or enter the vasculature (3, 4), whereas osteomyelitis, septic arthritis, infective endocarditis and pneumonia are deep-seated and systemic (5–13).

Introduction of penicillin to treat *S. aureus* bacteraemia patients in the early 1940s was immediately followed by isolation of penicillin resistant *S. aureus* strains (14). In *S. aureus*, penicillin resistance is mediated by the β-lactamase enzyme encoded by *blaZ*, which cleaves the β-lactam ring, thus disrupting the activity of the β-lactam antibiotic (14, 15). Methicillin, a penicillin derivative resistant to β-lactamase hydrolysis, was introduced in 1960s, but was quickly followed by the emergence of methicillin resistant *S. aureus* (MRSA) (16). Methicillin resistance was driven to the acquisition of the *mecA* gene on *Staphylococcus* cassette chromosome *mec* (SCC*mec*) elements, which encodes an alternative penicillin-binding protein, PBP2a, with a decreased affinity to β-lactams (17–21). In addition to *mecA*, auxiliary factors also contribute to high-level MRSA β-lactam resistance (22–36), including several involved in the synthesis of cell wall precursors, as well other physiological processes.

The ability of *S. aureus* to adapt to diverse host environments is linked to its ability to obtain essential nutrients from host tissues (37, 38), which in turn is dependent on metabolic reprogramming. A growing body of literature links central metabolic pathways to the pathogenicity of *S. aureus*, from its capacity to proliferate within the host, to the control of antibiotic resistance (22, 37–41). Thus, the identification of new drug targets and antibacterial strategies is reliant on first understanding virulence mechanisms associated with reprogramming of central metabolic pathways and their role in pathogenesis and antimicrobial resistance.

Bacteria synthesize macromolecules from 13 biosynthetic intermediates derived from glycolysis, the pentose phosphate pathway (PPP) and the tricarboxylic acid (TCA) cycle (42). *S. aureus* has the complete enzyme set for all three pathways, although it lacks a glyoxylate shunt (42). In addition to producing pentose precursors for biosynthesis of nucleotides and several amino acids, the PPP plays a critical role in cellular metabolism, maintaining carbon homeostasis by glucose turnover and contributing to the regeneration of reducing power in the form of NADPH (43–48). There are two branches in the PPP: the oxidative branch contributes to oxidative stress tolerance by generating reducing power in the form of NADPH/H^+^, and the non-oxidative branch produces ribose-5-P used in the *de novo* purine synthesis and the generation of nucleotide pools (ATP, ADP, AMP, c-di-AMP, GTP, GDP, GMP, ppGpp, pppGpp, IMP, XMP, etc.) for repair and synthesis of aromatic amino acids and peptidoglycan (47, 48). PPP activity is increased by environmental stress in Gram-positive organisms (48, 49).

Even though the contribution of glycolysis/gluconeogenesis and the PPP to intracellular persistence of *S. aureus* has been the subject of numerous studies (37, 38, 40, 45, 46, 48, 49), the role of these major glucose metabolism pathways in the antibiotic resistance of *S. aureus* remains largely unstudied. Mutations in PPP enzymes have been previously identified in slow growing-vancomycin intermediate *S. aureus* isolates (50).

We and others have previously reported that purine nucleotide homeostasis plays a key role in the regulation of β-lactam resistance in MRSA (49–53). Mutations in the *pur* operon and purine salvage pathway were associated with increased resistance, whereas exposure of MRSA to the purine nucleosides guanosine or xanthosine reduced β-lactam resistance (53). The purine-derived second messenger signalling molecules (p)ppGpp and c-di-AMP regulate β-lactam resistance, and exposure to exogenous guanosine downregulated c-di-AMP levels in *S. aureus* (53).

In this study, we investigated if mutations upstream of purine biosynthesis also control β-lactam resistance focusing on *pgl,* which is the only mutable gene in the oxidative phase of the PPP. We show that a *pgl* mutation in MRSA strain JE2, which leads to a slight growth defect in laboratory growth media, increased β-lactam resistance, but did not cause changes in PBP2a levels or peptidoglycan architecture. Carbon tracing and metabolomics experiments revealed increased flux to glycolysis and several cell envelope precursors. The susceptibility of wild-type JE2 to β-lactam antibiotics was dramatically increased in chemically defined medium containing glucose (CDMG), and accompanied by extensive cell lysis, whereas the *pgl* mutant remained highly resistant, exhibited a thick cell wall, intact septa and had a ruffled cell surface. Wheat germ agglutinin (WGA) binding assays indicated that wall teichoic acid (WTA)/lipoteichoic acid (LTA) levels were reduced or their composition altered in the *pgl* mutant. WTAs/LTAs and β-lactam resistance in the *pgl* mutant reverted to wild-type levels by mutations in the ABC transporter VraGF and cognate two-component regulatory system GraRS. These data reveal that metabolic reprogramming in an MRSA *pgl* mutant increases β-lactam resistance via VraFG/GraRS-dependent changes in cell envelope biogenesis.

## Results

### β-lactam resistance is increased in a MRSA *pgl* mutant

Extrapolating from previous data showing that purine metabolism controls β-lactam resistance (26, 41, 53–56), we turned our attention to the PPP, which produces ribose-5-P, a major substrate for purine and pyrimidine biosynthesis (Fig. 1). Given the important role of the PPP in central metabolism and production of reducing power, it is perhaps not surprising that mutations in the key enzymes in this pathway, including *zwf* and *gnd*, are not available in the Nebraska Transposon Mutant library (NTML) (57). However, the NTML does contain a mutation in the monocistronic *pgl* gene (SAUSA300_1902, NE202), which encodes 6-phosphogluconolactonase, the second enzyme in the oxidative phase of the PPP that converts 6-P-gluconolactone to gluconate-6-P.

**Fig. 1.**
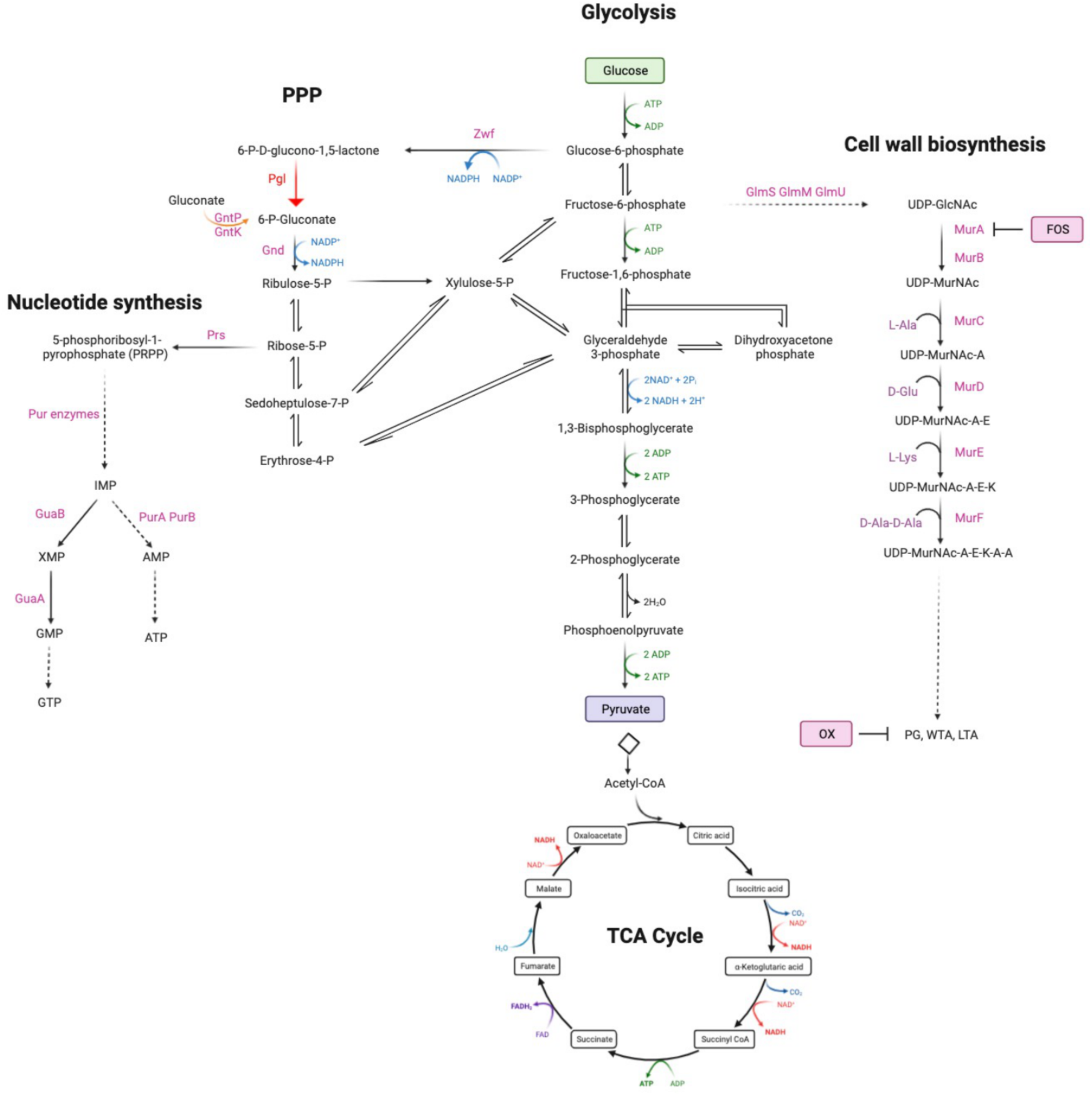
Summary of the oxidative phase of the pentose phosphate pathway including 6-phosphogluconolactonase (Pgl), that converts 6-P-gluconololactone to gluconate-6-P. For reference, key glycolysis, TCA cycle, nucleotide and cell wall biosynthetic pathway intermediates are also shown. Fructose-6-P is fluxed from glycolysis to peptidoglycan (PG), wall teichoic acid (WTA) and lipoteichoic acid (LTA) via UDP-N-acetylglucosamine (UDP-GlcNAc) and UDP-N-acetylmuramic acid (UDP-MurNAc). Fosfomycin (FOS) targets MurA which together with MurB is required for the conversion of UDP-GlcNAc to UDP-MurNAc. Oxacillin (OX) targets the transpeptidase activity of the penicillin binding proteins required for PG crosslinking. The putative gluconate shunt involves the export of 6-phosphogluconolactone, which spontaneously degrades to gluconate before being transported into the cell by the gluconate permease GntP and phosphorylated by the gluconate kinase GntK. Schematic made using Biorender.com.

When grown in Mueller Hinton 2% NaCl broth (MHB) the *pgl* mutant NE202 exhibited significantly increased resistance to cefoxitin in disk diffusion assays (zone diameters were 11mm for JE2 versus 8mm for *pgl*) and oxacillin (OX) in broth dilution assays (Table 1). Comparative whole genome sequencing analysis confirmed the absence of unexpected secondary mutations outside the *pgl* locus in NE202. The NE202 phenotype was verified by (i) showing that increased OX resistance was acquired by wild-type following transduction of the *pgl*::Erm^r^ allele and (ii) complementation of NE202 with the wild-type *pgl* gene (*pgl*_comp_) (Table 1). A *pgl*/*mecA* mutant was OX susceptible (Table 1) and Western immunoblotting revealed no differences in PBP2a expression between wild-type JE2, *pgl* and *pgl*_comp_ grown in TSB supplemented with OX 0.5 μg/ml (Fig. S1). Thus, high-level OX resistance in *pgl* was dependent on *mecA* but was not associated with increased PBP2a expression.

**Table 1.**
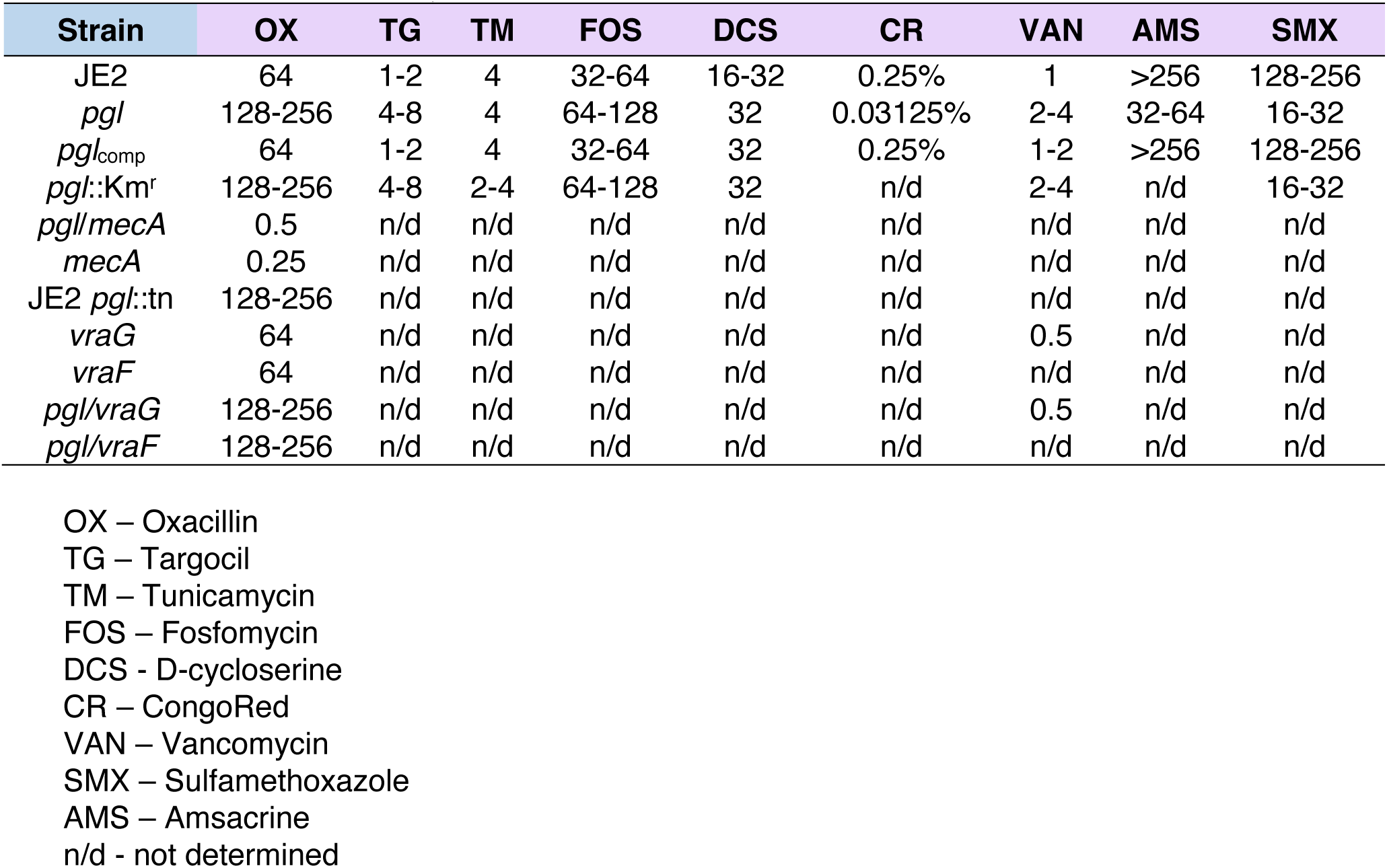
Minimum inhibitory concentrations (μg/ml) of strains used in this study to oxacillin (OX), targocil (TG), tunicamycin (TM), fosfomycin (FOS), D-cycloserine (DCS), Congo Red (CR), vancomycin (VAN), amacrine (AMS) and sulfamethoxazole (SMX) in Mueller Hinton Broth (+ 2% NaCl for OX); μg/ml

### The *pgl* OX resistance phenotype is glucose-regulated and independent of changes in peptidoglycan (PG) structure

Colonies of *pgl* were smaller than JE2 on MHA plates (Fig. S2A) and, in the absence of antibiotics, the *pgl* mutation negatively impacted growth in MHB (Fig. S2B), but to a lesser extent in LB, TSB and BHI (Fig. S2C-E). A *pgl* growth defect was also measured in chemically defined media with glucose (CDMG), but not in CDM without glucose (Fig. S2F,G). Growth of the complemented *pgl*_comp_ mutant was indistinguishable from the wild-type JE2 under all culture conditions tested (Fig. S2B-F). The mild growth defects of *pgl* in MHB and CDMG correlated with significantly increased OX MICs (Table 1, Fig. 2A), whereas the MIC of *pgl* in CDM (32-64 μg/ml) was more similar to wild-type JE2 (16-32 μg/ml; Fig. 2A). Notably, not only was *pgl* more resistant than wild-type JE2 in CMDG, but wild-type JE2 OX resistance was significantly reduced in this growth medium (MIC = 0.5 - 1 μg/ml; Fig. 2A). Wild-type JE2 and *pgl* grew similarly in CDM OX 10 μg/ml (Fig. 2B), whereas only *pgl* was able to grow in CDMG OX 10 μg/ml (Fig. 2C), revealing that this phenotype is glucose-regulated. The *pgl* mutation increased sensitivity to oxidative stress (H_2_O_2_) in CDMG (Fig. S3), similar to previous observations in *Listeria monocytogenes* using BHI media (58).

**Fig. 2.**
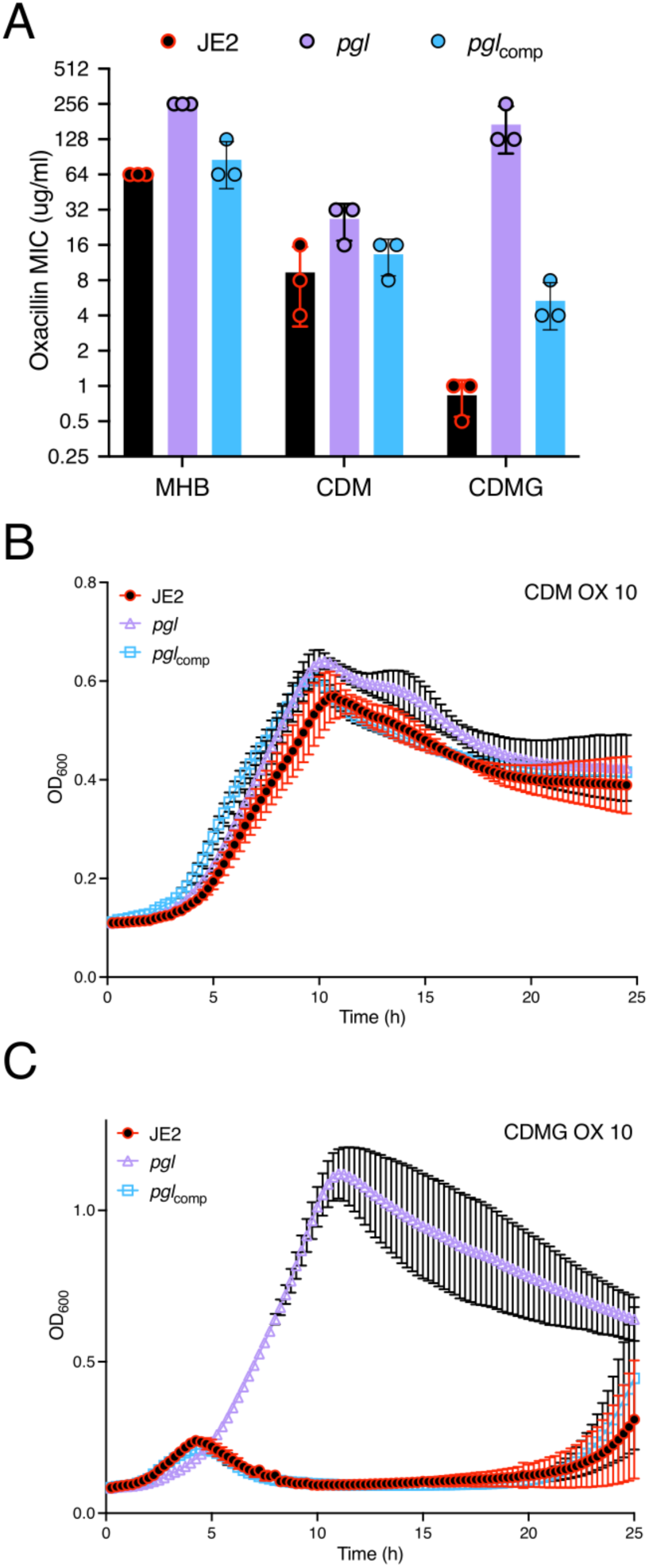
Mutation of *pgl* increases resistance to oxacillin. **A.** Oxacillin MICs of JE2, *pgl* and the complemented *pgl* mutant in Mueller Hinton broth with 2% NaCl (MHB), chemically defined media (CDM) and CDM with glucose (CDMG). **B and C.** Growth of JE2, *pgl* and *pgl*_comp_ for 25 hrs at 35°C in CDM (B) and CDMG (C) supplemented with OX 10 μg/ml. Growth (OD_600_) was measured at 15 min intervals in a Tecan plate reader. Data are the average of 3 independent experiments and error bars represent standard deviation.

Confocal microscopy revealed that the diameter of *pgl* cells from overnight CDMG grown cultures was significantly smaller than wild-type JE2 or *pgl*_comp_ cells (Fig. 3A,B). The OX-induced increase in MRSA cell size, which we and others have previously reported (31, 53, 59–61), was more significant in wild-type JE2 and *pgl*_comp_ than the *pgl* mutant (Fig. 3C). Furthermore, the increased cell size of wild-type JE2 and *pgl*_comp_ in CDMG OX was associated with a dramatic increase in the number of cells undergoing visible lysis (Fig. 3D), an observation consistent with the abrupt decline in the OD_600_ of wild-type JE2 and *pgl*_comp_ cultures after 4-5 h growth under these growth conditions (Fig. 2C). Quantitative PG compositional analysis of muramidase-digested muropeptide fragments revealed similar oligomerisation profiles and crosslinking for wild-type JE2, *pgl* and the *pgl*_comp_ strains grown in CDMG, or CDMG supplemented with sub-inhibitory 0.05 μg/ml OX, MHB 2% NaCl, MHB 2% NaCl supplemented with 0.5 μg/ml OX, (Fig. S4A-D) The total PG content was also similar for all three strains under these growth conditions (Fig. S4E-H). Thus, in addition to the unchanged PBP2a expression (Fig. S1), increased *pgl* OX resistance was unrelated to changes in PG structure or amount (Fig S4).

**Fig. 3.**
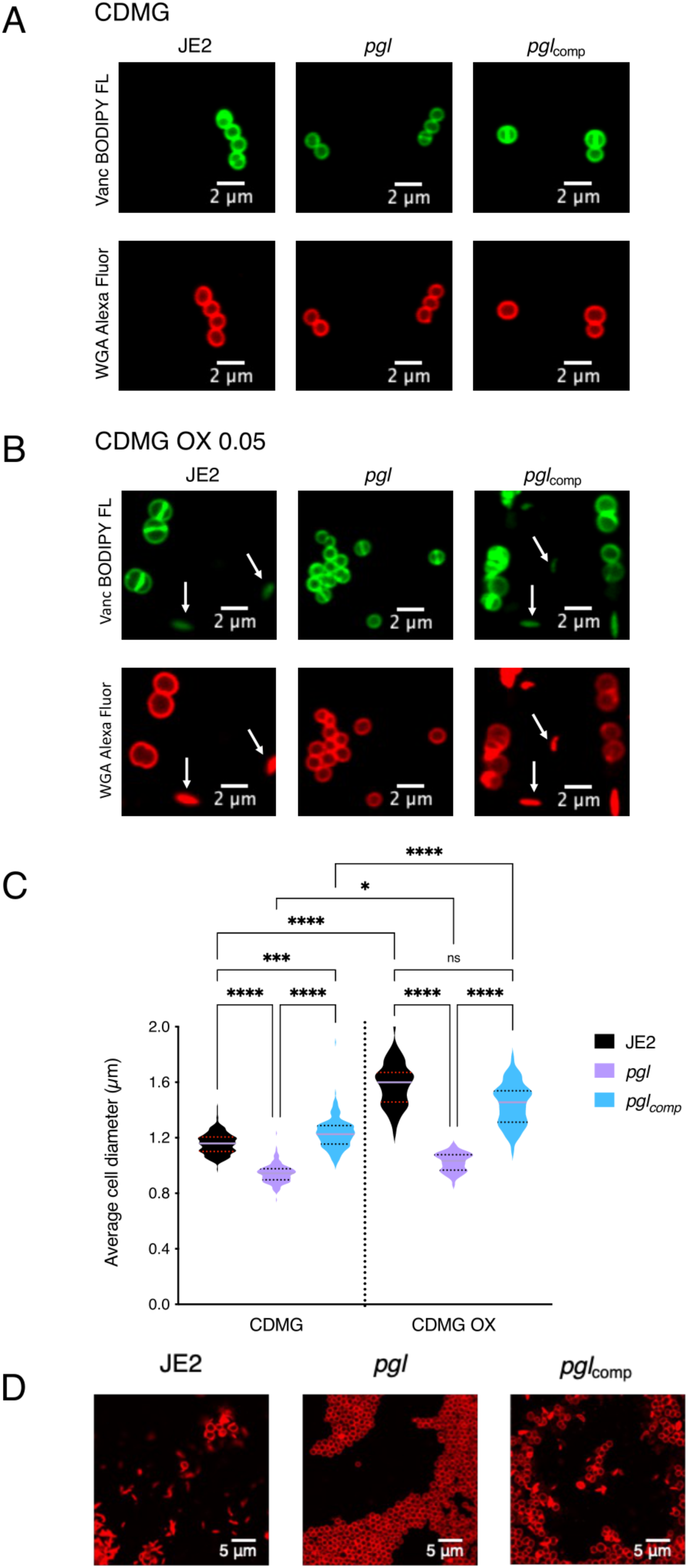
Mutation of *pgl* reduces cell size and prevents OX-induced cell lysis in CDMG. **A and B.** Representative microscopic images of JE2, *pgl* and *pgl*_comp_ cells grown in CDMG (A) or CDMG supplemented with OX 0.05 μg/ml (B) and labelled with vancomycin BODIPY FL, which binds to the terminal D-ala-D-ala in the peptidoglycan stem peptide (green, top panel) or WGA Alexa Fluor 594, which binds to GlcNAc and other sugars in the cell envelope (red, bottom panel). **C.** Average diameter of JE2, *pgl* and *pgl*_comp_ cells grown in CDMG or CDMG OX. Images of cells from three biological replicates were acquired using Fv3000 confocal microscope and software, 50 cells measured per biological replicate (200 cells in total) for CDMG and 60 cells in total counted for CDMG OX (due to cell lysis), and the violin plots for the four biological replicates were generated using GraphPad Prism V9. Asterisks indicate statistically significant difference according to using a Kruskal-Wallis test followed by a Dunn’s multiple comparison test. Adjusted p-values * p<0.05, *** p<0.001 and **** p<0.0001 are indicated. **D.** Extensive lysis of JE2 and *pgl*_comp_ (but not *pgl*) in CDMG OX 0.05 μg/ml cultures. Cells were labelled with WGA Alexa Fluor 594 and representative microscopic images are shown.

### Exogenous D-gluconate or the *gntPK* gluconate shunt genes do not play a role in the *pgl* OX resistance phenotype

In *Escherichia coli* and *L. monocytogenes*, 6-phosphogluconolactone that accumulates in *pgl* mutants is dephosphorylated to labile gluconolactone, which is exported out of the cell where it spontaneously hydrolyses to gluconate (58, 62). In *S. aureus*, the predicted gluconate shunt genes *gntP* (SAUSA300_2442) and *gntK* (SAUSA300_2443) are co-located on the chromosome with the *gntR* regulator. In a previous RNAseq analysis, we reported that *gntP* was upregulated by OX (63). Growth of wild-type JE2, *pgl* and *pgl*_comp_ in CDMG supplemented with 5 g/l D-gluconate alone or with 10 μg/ml OX was similar (Fig. S5A, B). Furthermore, inactivation of *gntP* or *gntK* in the *pgl* mutant did not affect OX resistance (Fig. S5C). Therefore, exogenous D-gluconate does not regulate OX resistance under conditions tested, and the gluconate shunt genes are not required for the viability or increased OX resistance of the *pgl* mutant.

### Inactivation of *pgl* reduces carbon flux through PPP

Liquid chromatography-tandem mass spectrometry analysis was used to trace [1,2-^13^C_2_] glucose flux through glycolysis and the PPP in wild-type JE2, *pgl* and *pgl*_comp_. As described previously (64), six-carbon [1,2-^13^C] glucose can be metabolised via glycolysis and the PPP to produce three-carbon ^13^C_2_-pyruvate (M+2) and ^13^C_1_-pyruvate (M+1), respectively (Fig. 4A). The M+1 fraction is produced following a decarboxylation reaction in the PPP that releases ^13^CO_2_ (Fig. 4A). M+1 pyruvate levels were reduced in *pgl*, indicative of reduced PPP activity (Fig. 4B), whereas M+2 pyruvate levels derived primarily from glycolysis, were similar (Fig. 4B). The M+2/M+1 ratio further illustrated the impaired PPP activity of *pgl* and showed >2-times more pyruvate generated directly from glucose entering glycolysis in *pgl* than in wild-type JE2 or *pgl*_comp_ (Fig. 4C).

**Fig. 4.**
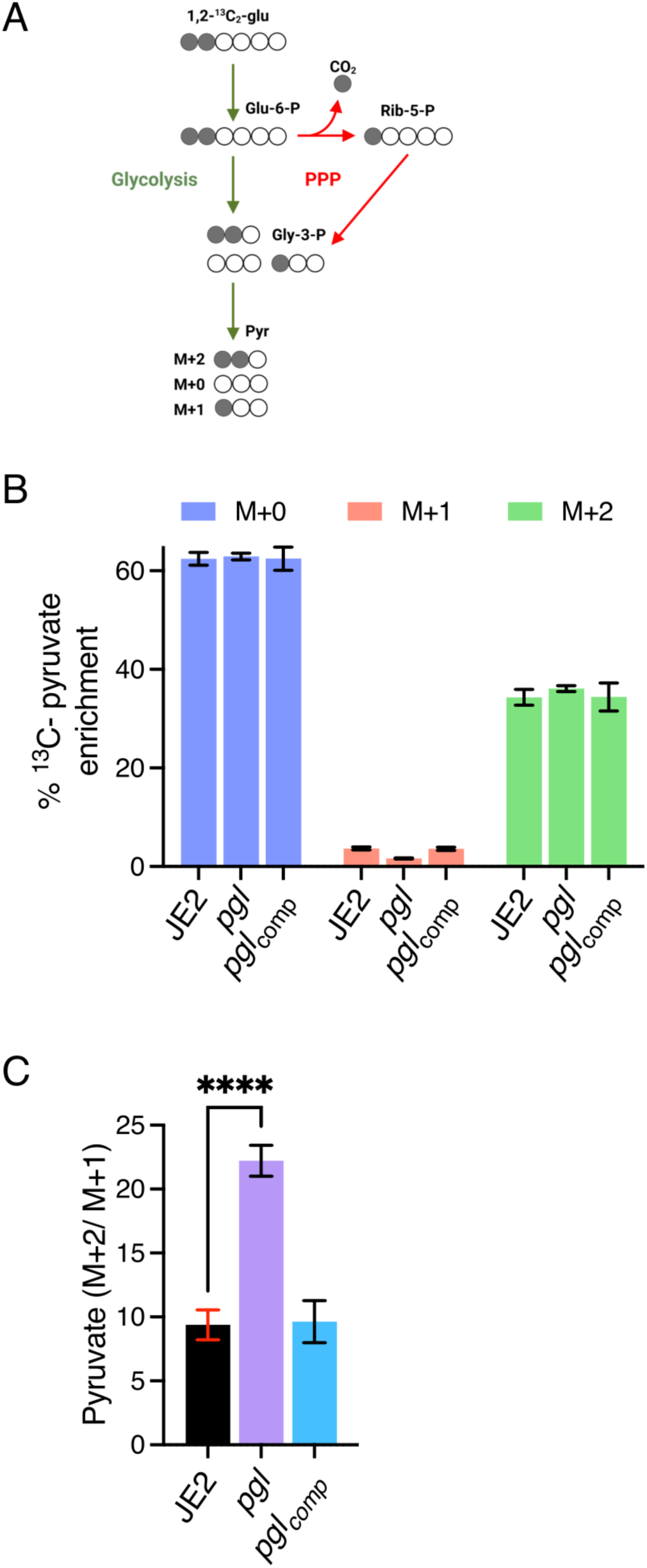
PPP activity is impaired in the *pgl* mutant. **A.** JE2, *pgl* and the complemented *pgl* mutant were grown in CDM [1,2-^13^C]Glucose and fluxes via glycolysis and the pentose phosphate pathway (PPP) were compared as described previously (64). The M+2 pyruvate is unique to glycolysis and the M+1 pyruvate to PPP. Thus, the M+2/M+1 ratio is indicative of carbon flux through glycolysis relative to PPP. The M+0 pyruvate can arise from different sources including the unlabeled part of the [1,2-^13^C]Glucose and pyruvogenic amino acids that are consumed alongside glucose. **B.** Relative levels of M+1 pyruvate indicative of PPP activity and M+2 pyruvate indicative of glycolytic activity in JE2, *pgl* and *pgl*_comp_. **C.** The M+2/M+1 ratio indicative of pyruvate produced directly from glucose flux through glycolysis in JE2, *pgl* and *pgl*_comp_. Data are the average of three independent experiments and standard deviations are shown. Significant differences were determined using ordinary one-way ANOVA with Dunnett’s multiple comparison using GraphPad Prism V9 and adjusted p-value **** p<0.0001 is indicated.

### OX resistance in the *pgl* mutant is independent of the TCA cycle or glucogenic and ketogenic amino acids

HPLC was used to investigate if redirected glucose flux from the PPP to glycolysis impacted consumption of amino acids in CDMG, and revealed that levels of threonine, the branched chain amino acids (BCAAs) valine, leucine and isoleucine, as well as phenylalanine, tryptophan and tyrosine, histidine, methionine and aspartic acid were increased in the supernatant of *pgl* cultures compared to JE2 or *pgl*_comp_ after 7.5 h growth (Fig. S6A). Interestingly, the levels of the TCA cycle intermediates malate, succinate and particularly α-ketoglutarate were also increased in CDMG supernatants of *pgl* (Fig. S6B), which may be consistent with a reduced requirement for glucogenic and ketogenic amino acids. To investigate this proline dehydrogenase (*putA*::Em^r^), isopropylmalate dehydrogenase (*leuB*::Em^r^) and glutamate dehydrogenase (*gudB*::Em^r^) mutations, predicted to interfere the flux of amino acids to α-ketoglutarate and/or pyruvate, were transduced from the NTML (57) into *pgl*::Km^r^. Growth of the resulting *pgl*/*putA*, *pgl*/*leuB,* and *pgl*/*gudB* mutants in CDMG and CDMG OX was similar to *pgl*::Km^r^ (Fig. S6C,D). Similarly the *pgl* TCA cycle double mutants *pgl*/*sdhA*, *pgl*/*sucA* and *pgl*/*sucC* remained capable of growing in CDMG OX (Fig. S6C,D), although *pgl*/*sucC* exhibited an extended lag phase in keeping with our previous report that *sucC* mutation re-sensitizes MRSA to β-lactam antibiotics due to increased accumulation of succinyl CoA (39). Collectively, these data indicate that an intact TCA cycle or the accumulation of TCA cycle intermediates and glucogenic/ketogenic amino acids in culture supernatants was not associated with the increased β-lactam resistance of the *pgl* mutant.

### Increased resistance to β-lactam antibiotics in *pgl* is promoted by redirected carbon flux to cell wall precursors

Whole cell metabolomics was performed on JE2, *pgl* and *pgl*_comp_ grown in CDMG or CDMG OX (Fig. 5). Consistent with the important role of the PPP in the generation of reducing power and nucleotide biosynthesis, levels of key redox carriers and six nucleotides were significantly reduced in *pgl* and restored to JE2 levels in the complemented mutant (Fig. 5). Interestingly, reduced nucleotide levels correlated with a 2-4-fold increase in the susceptibility of *pgl* mutant to sulfamethoxazole, which inhibits dihydropteroate synthetase in the folate synthesis pathway (Table 1). Levels of sedoheptulose 7-P which is downstream of Pgl in the PPP was also reduced in *pgl*, reaching significance in CDMG, whereas ribose 5-P and erythrose 5-P were significantly increased (Fig. 5), indicative of complex metabolic reprogramming in the *pgl* mutant.

**Fig. 5.**
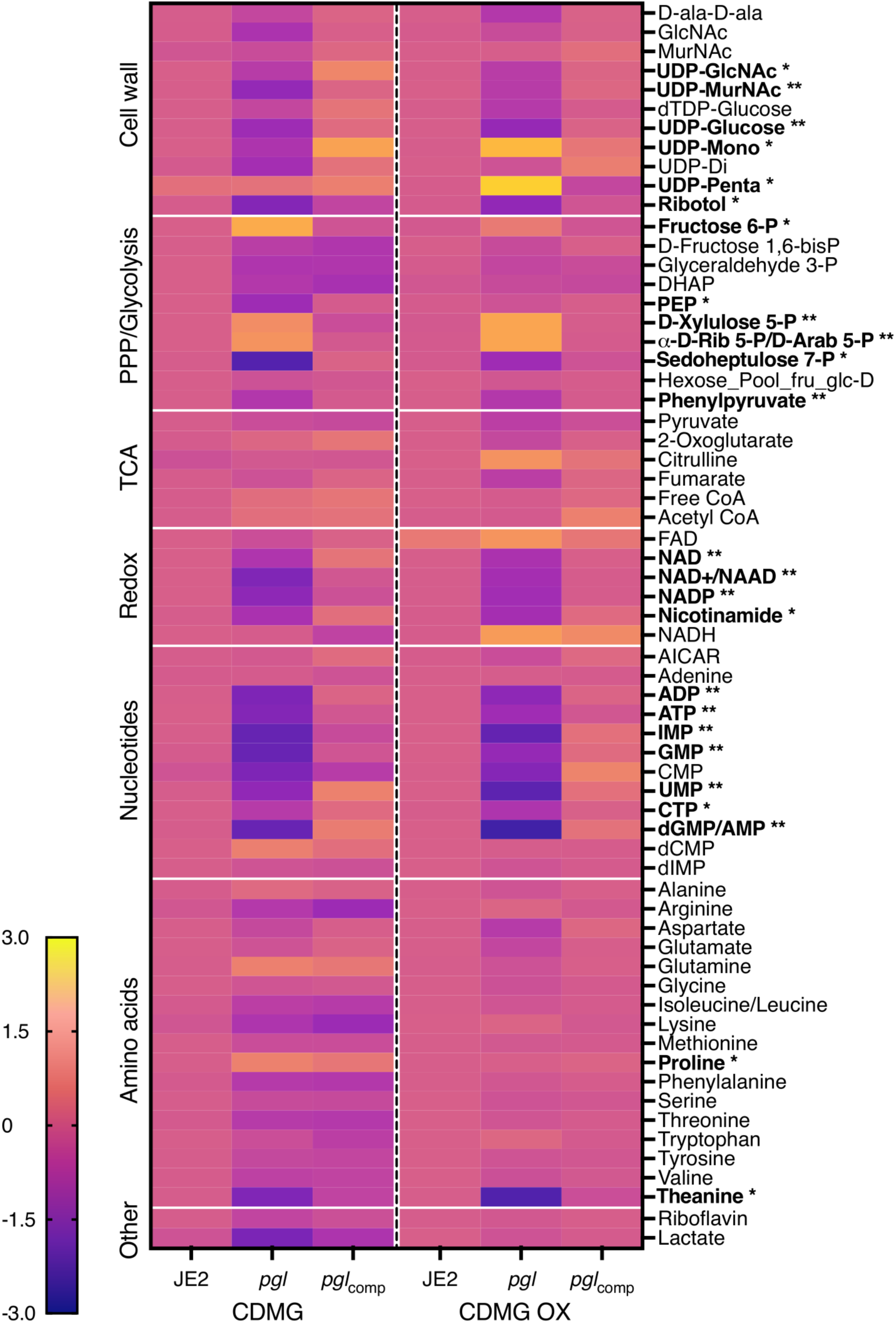
Heatmap comparison of cell wall, pentose phosphate pathway (PPP)/glycolysis, TCA cycle, redox, nucleotides and amino acid metabolites in JE2, *pgl* and *pgl*_comp_. Whole cell metabolomics was performed on JE2, *pgl* and *pgl*_comp_ grown in CDMG and CDMG OX 10 μg/ml. Data presented are the average of three biological replicates (2 biological replicates for FAD) analysed using GraphPad Prism V9. Individual metabolite levels that were significantly different using a one-way ANOVA with Turkey’s post-hoc in *pgl* grown in CDMG, CDMG OX or both are highlighted in bold text. * significant difference in either CDMG or CDMG OX. ** significant difference in both CDMG and CDMG OX.

Consistent with the [1,2-^13^C] glucose tracing experiments, accumulation of fructose 6-P from which cell wall precursors are derived, was increased in CDMG OX and significantly increased in CDMG (Fig. 5). Furthermore, the downstream glycolytic intermediates fructose 1,6-bis-P, dihydroxyacetone phosphate (DHAP), glyceraldehyde 3-P and phosphoenolpyruvate (PEP) were reduced (Fig. 5). Although there are several possible explanations for this, one possibility is that the accumulated fructose 6-P may be fluxed to the PPP or cell wall. Indeed, significantly increased levels of UDP-mono and UDP-penta were measured in *pgl* grown in CDMG OX, but not in CDMG (Fig. 5). In contrast, the levels of UDP-GlcNAc and UDP-MurNAc were significantly decreased (Fig. 5), perhaps reflecting increased consumption of these substrates in the production of UDP-mono and UDP-penta in CDMG OX. Increased accumulation of UDP-mono and UPD-penta correlated with the increased resistance of the *pgl* mutant to fosfomycin (FOS) (Table 1, Fig. S7), an antibiotic that inhibits the MurA enzyme, which together with MurB catalyses the conversion of UDP-GlcNAc to UDP-MurNAc (Fig. 1). Furthermore, *pgl* exhibited significantly increased resistance to an OX/FOS combination compared to wild-type JE2 in a checkerboard dilution assay (Fig. S7). Broth microdilution susceptibility experiments revealed that the *pgl* mutant was 1-2-fold more resistant to vancomycin (VAN), which targets the terminal D-ala-D-ala of the PG stem peptide (Table 1).

Taken together, these data indicate that redirected carbon flux to cell wall precursors in *pgl* contributes to the increased resistance to β-lactam antibiotics. Furthermore, *pgl* viability appears to be underpinned by a complex and regulated interconversion of glycolytic and PPP intermediates, which may also explain why the glycolytic shunt genes are dispensable for the growth of the *pgl* mutant under these culture conditions.

### Mutation of *pgl* alters susceptibility to antimicrobial agents targeting wall teichoic acids (WTAs) and lipoteichoic acids (LTAs) and is accompanied by morphological changes in the cell envelope

The MICs of wild-type JE2 and *pgl* to the TarO inhibitor tunicamycin were the same, whereas *pgl* was more resistant to the TarGH inhibitor targocil and more susceptible to the D-alanylation inhibitor amsacrine (Table 1), revealing different effects of antimicrobial agents targeting distinct steps in WTA biosynthesis. TarO catalyzes the transfer of N-acetylglucosamine-1-phosphate from UDP-GlcNAc to undecaprenyl-P to initiate WTA synthesis (65). The TarGH ABC transporter transports WTAs across the cytoplasmic membrane (66), and the polymer is then D-alanylated by the DltABCD complex (67). The *pgl* mutant was also more sensitive to Congo red which targets the LTA synthase LtaS (68) (Table 1). Importantly LTA is also D-alanylated by DltABCD. The susceptibility of *pgl* to D-cycloserine, which targets the alanine racemase and ligase enzymes in the D-ala-D-ala pathway was unchanged when compared to wild-type, and the metabolomic analysis also showed no significant differences in the levels of D-ala-D-ala in wild-type JE2, *pgl* and *pgl*_comp_ (Fig. 5).

Transmission electron microscopy (TEM) revealed that *pgl* cells grown in CDMG OX had visibly ruffled surface characteristics, and thick, intact septa compared to JE2 cells (Fig. 6). Consistent with previous microscopic analysis (Fig. 3), TEM revealed defective/truncated septa in dividing wild-type cells, as well as cells undergoing lysis (Fig. 6). In contrast wild-type and *pgl* cells grown in the absence of OX were largely similar (Fig. S8). Taken together these data suggest that cell envelope changes in the *pgl* mutant are the result of altered activity of the TarGH, LtaAS-YpfP and DltABCD membrane complexes involved in export and D-alanylation of WTAs and LTAs that collectively contribute to increased OX resistance.

**Fig. 6.**
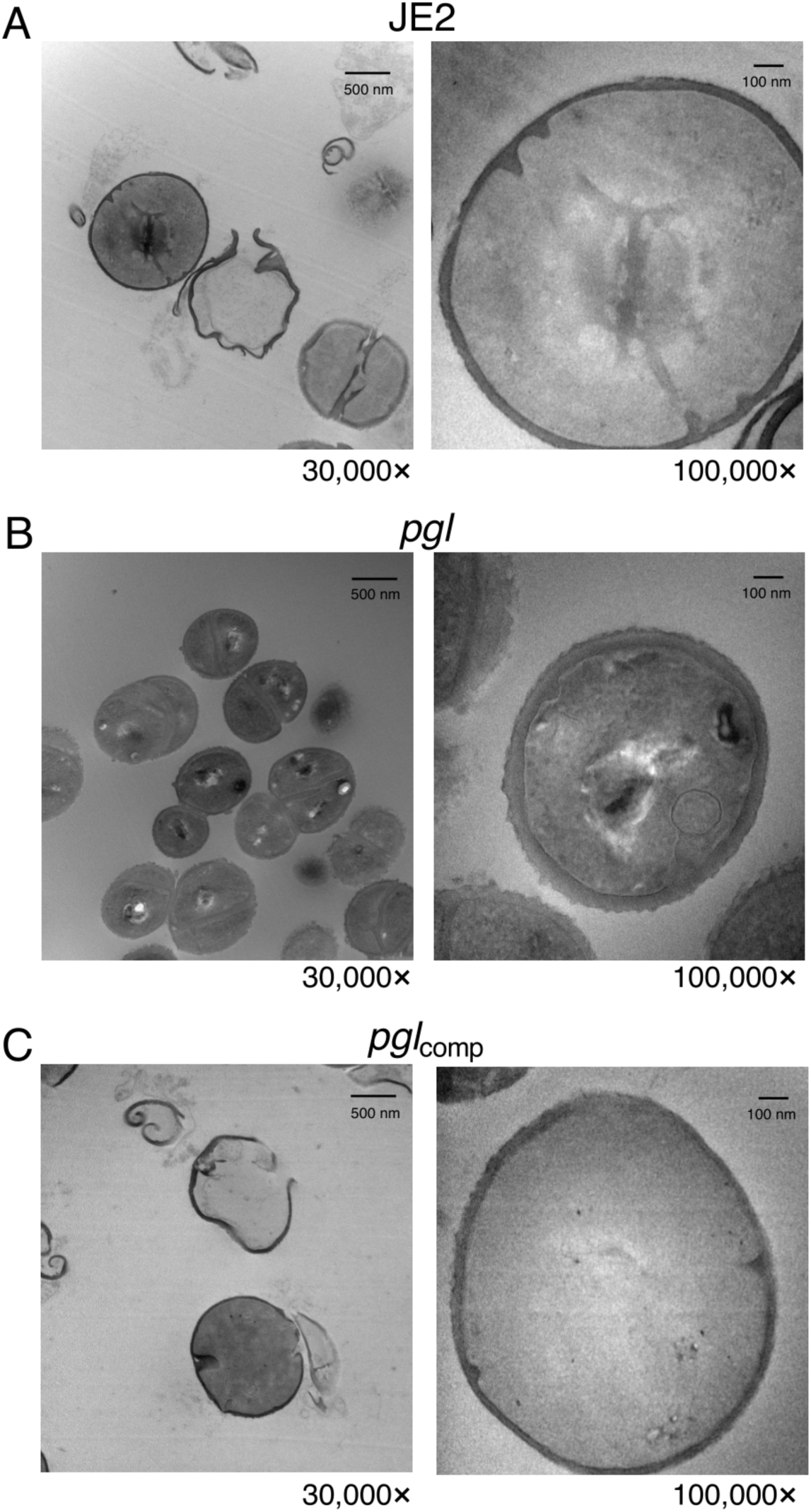
Increased OX resistance in the *pgl* mutant is associated with a ruffled surface morphology, a thicker cell wall and thicker septa between dividing cells. Transmission electron microscopy at 30,000× (left) and 100,000× (right) magnification was performed on JE2 (A), *pgl* (B) and *pgl*_comp_ (C) cells collected from exponential phase cultures grown for 4.5 h in CDMG OX 1 μg/ml normalized to OD_600_ = 1 in PBS before being fixed and thin sections prepared. Representative cells from each strain are shown. Scale bars represent 500 nm at 30,000× or 100 nm at 100,000× magnification.

### OX resistance in the *pgl* mutant is dependent on *vraF* and *vraG*

During experiments to remove the Erm^r^ marker in NE202 a *pgl* markerless transposon mutant that had reverted to wild-type patterns of growth in CDMG OX was isolated (Fig. 7A, B). Whole genome sequencing of this mutant, designated *pgl*R1, identified a thymine deletion 73bp upstream of *putA* and a Gln_394_STOP substitution in VraG. Construction of *pgl* double and triple mutants revealed that the *pgl* OX resistance phenotype was dependent on *vraG* and not *putA* (Fig. 7A, B). (Fig. 7A, B). *vraG* encodes a membrane permease, which together with its cognate ATPase, VraF, comprises an ABC efflux pump previously implicated in resistance to cationic antimicrobial peptides, polymyxin B and vancomycin (69–73), potentially via the export of cell wall/teichoic acid precursors or modifying subunits (71). Consistent with this, a *pgl*/*vraF* mutant grew similarly to *pgl*/*vraG* and JE2 in CDMG and CDMG OX (Fig. 7A, B). VraFG is also part of a multicomponent complex with the glycopeptide resistance-associated GraRS two component system, that regulates *vraFG* as well as the *dltABCD* operon and *mprF* (69–71). A *pgl*/*graR* mutant exhibited an intermediate phenotype when grown in CDMG OX compared to JE2 and the *pgl*/*vraF* or *pgl*/*vraG* mutants (Fig. 7B) revealing a role for the entire VraFG/GraRS complex in the *pgl* OX resistance phenotype.

**Fig. 7.**
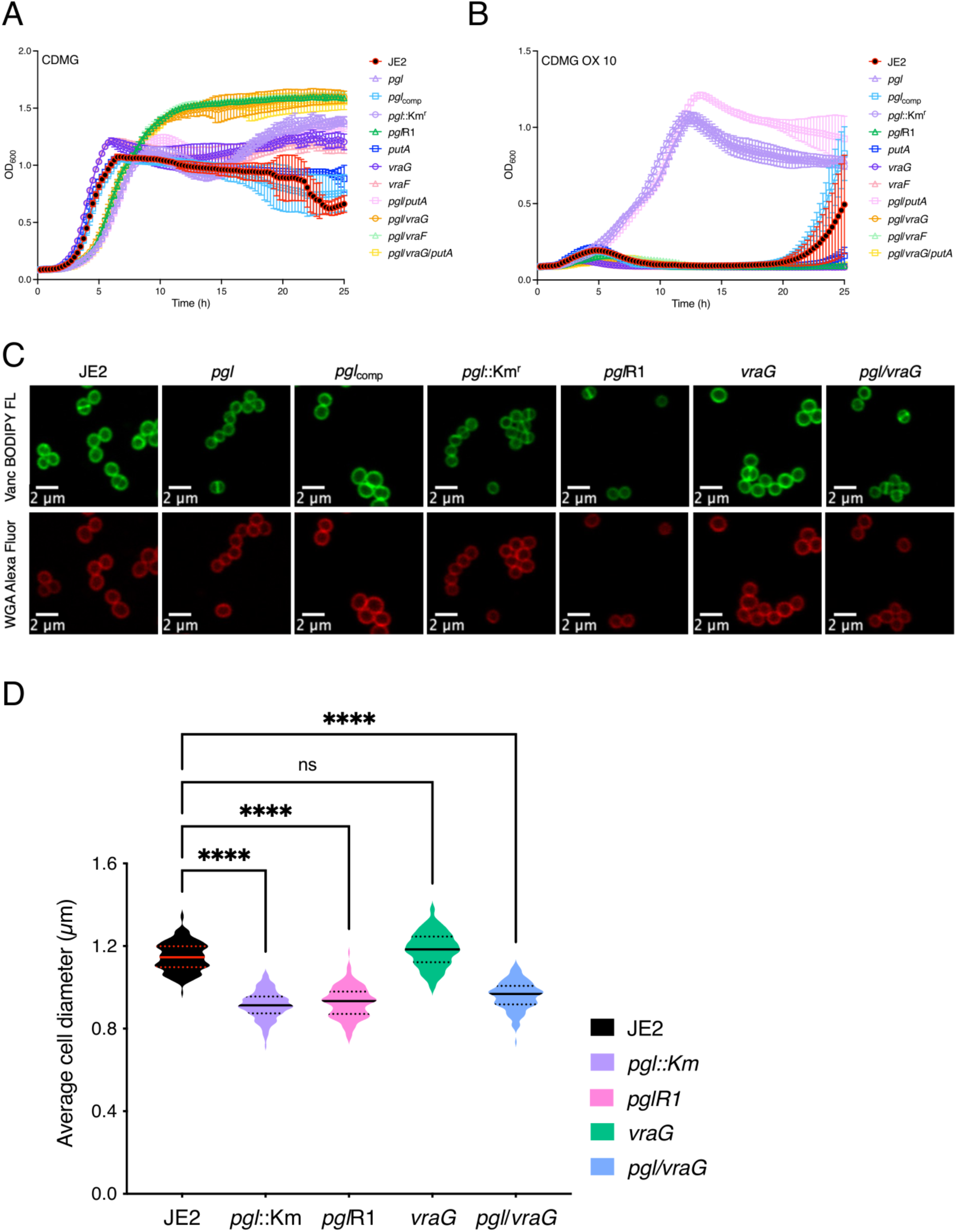
Mutation of *vraG* restores wild type OX resistance, but not cell size, in the *pgl* mutant grown in CDMG. A and B. Growth of JE2, *pgl*::Km^r^, *pgl*R1 *putA*, *vraG*, *pgl*/*putA*, *pgl*/*vraG*, *pgl*/*putA*/*vraG* for 25 hrs at 35°C in CDMG (A) and CDMG supplemented with OX 10 μg/ml (B). Growth (OD_600_) was measured at 15 min intervals in a Tecan plate reader. Data are the average of 3 independent experiments and error bars represent standard deviation. **C.** Representative microscopic images of JE2, *pgl*, *pgl*_comp_, *pgl*::Km^r^, *pgl*R1, *vraG* and *pgl/vraG* cells grown in CDMG and labelled with vancomycin BODIPY FL (green, top panel) or WGA Alexa Fluor 594 (red, bottom panel). **D.** Average diameter of JE2, *pgl*::Km^r^, *pgl*R1, *vraG* and *pgl*/*vraG* cells grown in CDMG. Images of cells from three biological replicates were acquired using Fv3000 confocal microscope and software, 50 cells measured per biological replicate (150 cells in total) and the violin plots for the three biological replicates were generated using GraphPad Prism V9. Asterisks indicate statistically significant difference according to using a Kruskal-Wallis test followed by a Dunn’s multiple comparison test. Adjusted p-values **** p<0.0001 or ns, not significant are indicated.

Interestingly, the increased VAN MIC of the *pgl* mutant grown in MHB was reduced from 2-4 μg/ml to 0.5 μg/ml in *pgl*/*vraG* (Table 1) further indicating a reversal of cell envelope changes. However, *pgl*/*vraG* and *pgl* cells were the same size in CDMG and CDMG OX (Fig. 7C, D) demonstrating that the *vraG* mutation does not reverse other central metabolism-related phenotypes in the *pgl* mutant.

In CDMG, the OX MICs of *putA*, *vraF*, *vraG* and *pgl*/*putA* were the same as JE2, whereas *pgl*/*graR* was reduced to 32-64 μg/ml and *pgl*/*putA*/*vraG*, *pgl*/*vraF* and *pgl*/*vraG* were reduced to 8 μg/ml compared to 128-256 μg/ml for *pgl* and *pgl*::Km^r^ (Fig. S9). Interestingly, in MHB 2% NaCl, the increased OX MIC of the *pgl* mutant (128-256 μg/ml) was not reversed in the *pgl*/*vraG* or *pgl*/*vraF* mutants (Table 1), underlining the importance of exogenous glucose in *pgl*-related phenotypes and indicating that VraFG-dependent OX resistance in the *pgl* mutant is environmentally regulated.

### Evidence that reduced levels of teichoic acids in the *pgl* mutant are restored by *vraF* or *vraG* mutations

To compare levels of teichoic acids in the *pgl*, *vraF*, *vraG* and *graR* strains, wheat germ agglutinin (WGA) Alexa Fluor 594 binding assays were performed using fluorescence microscopy. WGA binds to GlcNAc and other sugars in WTA, LTA and PG. WGA binding to *pgl* and *pgl*::Km^r^ cells was significantly reduced compared to the JE2, *vraF*, *vraG* and *graR* strains (Fig. 8). Furthermore, WGA binding was restored to wild-type levels in *pgl*R1, *pgl*/*vraF, pgl*/*vraG, pgl*/*putA*/*vraG* and *pgl*/*graR* (Fig. 8A). Increased Congo Red susceptibility of the *pgl* mutant (Table 1) (Fig. 8B), which is consistent with reduced WGA binding was also reversed in the *pgl*/*vraG* mutant and complemented, albeit partially in *pgl*_comp_ (Fig. 8B). Levels of ribitol, the backbone for WTAs, were also reduced in *pgl* grown in CDMG OX and significantly reduced in CDMG (Fig. 5).

**Fig. 8.**
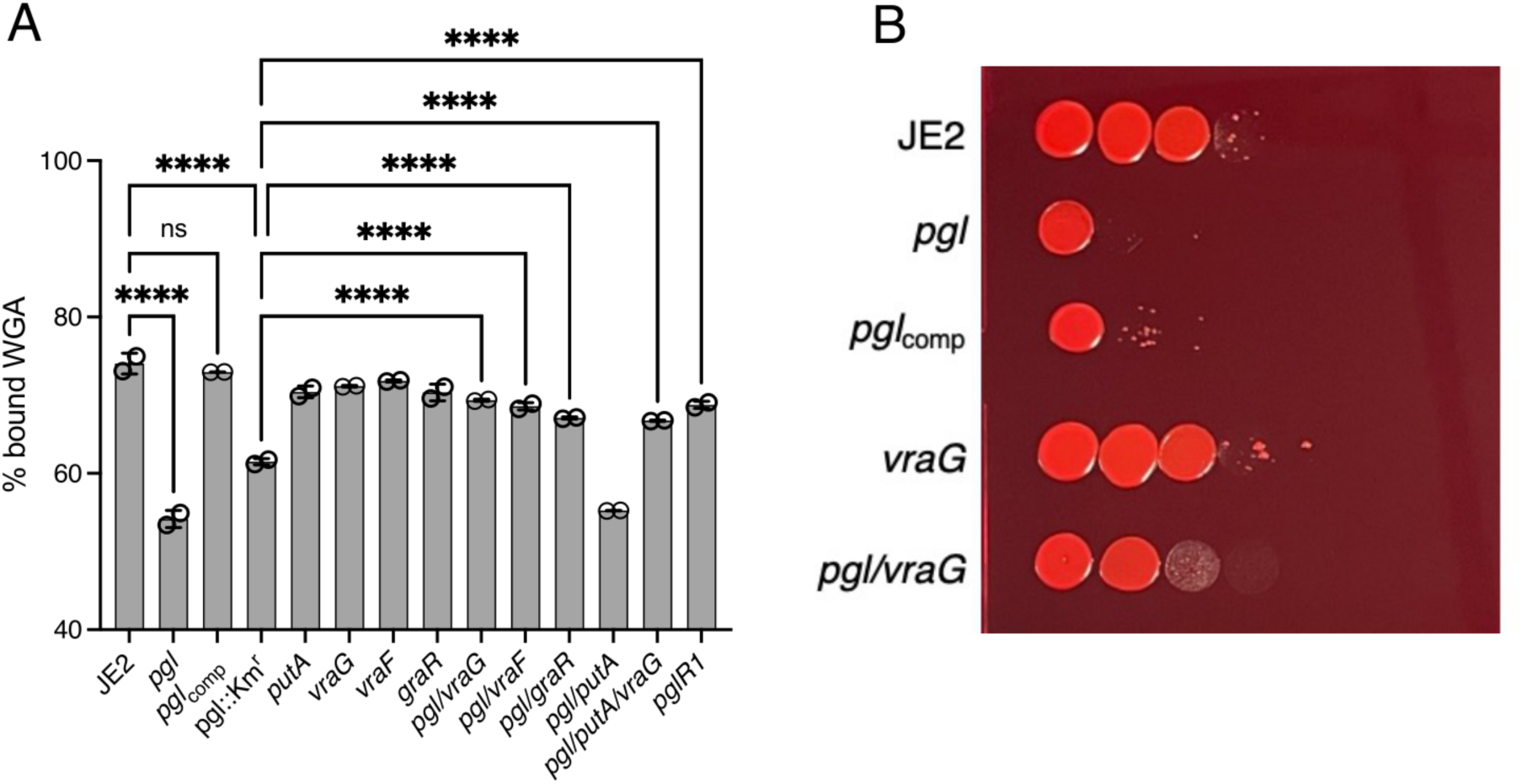
Evidence for reversal of teichoic acid-associated phenotypes in the *pgl* mutant by *vraF* and *vraG* mutations. **A.** Comparison of wheat germ agglutinin (WGA) Alexa Fluor 594 binding to JE2, *pgl*, *pgl*_comp_, *pgl*::Km^r^, *vraG, vraF, graR, putA*, *pgl*/*vraG, pgl/vraF, pgl/graR, pgl/putA, pgl/putA/vraG and pgl*R1 cells grown for 4.5 h in CDMG OX using fluorescence microscopy at 594nm excitation/618nm detection. The data are the average of 3 independent experiments and error bars represent standard deviation. Significant differences were determined using a two-way ANOVA with Turkey’s post-hoc analysis. Adjusted p-values **** p<0.0001 or ns, not significant are indicated. **B.** Mutation of *vraG* partially restores Congo red resistance in the *pgl* mutant. 10-fold serial dilutions of JE2, *pgl*, *pgl*_comp_, *vraG* and *pgl*/*vraG* inoculated onto TSA supplemented with 0.125% Congo red and grown for 24 h at 37°C. This experiment was repeated three times and a representative plate is shown.

Nevertheless, the correlation between reduced levels of teichoic acids and increased OX resistance in *pgl* is difficult to explain, perhaps suggesting that teichoic acid structures are different, as evidenced by the ruffled surface morphology. WTA glycosylation, which has previously been implicated in increased OX resistance (63, 74), was ruled out because the OX MICs of the *pgl* mutant was unaffected by mutations in the WTA α and β glycosylase genes *tarM* and *tarS* (Fig. S9). Future studies to characterise WTA/LTA polymer length and decoration (75–77), and patterns of LTA release (27, 78), which may be controlled by VraFG/GraRS will be informative. In summary, the data presented here reveal that extensive metabolic reprogramming in a MRSA *pgl* mutant is accompanied by increased OX resistance, which is associated with redirected carbon flux to cell envelope precursors and VraFG-GraRS-dependent resistance to OX-induced lysis.

## Discussion

Beyond the central role of the oxidative phase of the PPP in producing reducing power and 5 carbon sugars for nucleotide and amino acid biosynthesis, its contribution to other phenotypes in bacteria remains much less understood due to the essentiality of most enzymes in the pathway. One exception to this is 6-phosphogluconlactonase (Pgl). Here, for the first time, we report a role for *pgl* in the control of MRSA β-lactam antibiotic resistance, growth, cell size and cell surface morphology. Our analysis revealed pleiotropic effects of the *pgl* mutation on (i) the PPP itself and downstream nucleotide biosynthesis, (ii) glycolysis and the TCA cycle and (iii) flux to cell wall, WTA and LTA precursors. All three of these pathways have previously been implicated in the control of MRSA β-lactam resistance providing a multifaceted explanation for the OX resistance of the *pgl* mutant.

Although OX MICs of the wild-type JE2 and *pgl* mutant were dependent on the culture media, the *pgl* mutant was always significantly more resistant and the most striking difference between the two strains was measured in chemically defined media with glucose (CDMG), which is the substrate for the PPP. Strikingly, the wild-type JE2 OX MIC was reduced to 1 μg/ml in CDMG, compared to 64 μg/ml in MHB 2% NaCl, whereas the *pgl* OX MIC was similar in both culture media (128-256 μg/ml). Given that the JE2 OX MIC was 4-16 μg/ml in CDM, in which growth of *S. aureus* is dependent on amino acid catabolism, it appears that glucose plays a significant role in controlling OX susceptibility in JE2 but not in the *pgl* mutant in which central metabolism is significantly perturbed. MHB 2% NaCl is the standard culture medium used to measure the susceptibility of *S. aureus* clinical isolates to oxacillin in diagnostic laboratories, and these experiments raise the question of whether CDMG may be more physiologically relevant in terms of predicting the *in vivo* effectiveness of β-lactams in patients with MRSA infections.

The *pgl* gene has previously been mutated in *E. coli* and *L. monocytogenes* (58, 62), leading to an accumulation of gluconate, which can be transported back into the cell and phosphorylated, thus potentially bypassing the Pgl-catalysed reaction in the PPP (58, 62). However, in *S. aureus* the slower growth and OX resistance phenotypes of the *pgl* mutant were not dependent on the gluconate transport and catabolism genes *gntPK* or the presence of exogenous D-gluconate in the culture media. Thus, questions remain about the importance and regulation of the gluconate shunt in *S. aureus*.

In CDMG, *pgl* mutant cell size was significantly reduced compared to wild-type. Reduction in cell size may correlate with increased β-lactam resistance of *pgl*, as previously reported for a c-di-AMP phosphodiesterase *gdpP* mutant (41). In addition to the previously reported OX-induced increase in cell size (31, 53, 59, 61), a dramatic cell lysis phenotype was also observed in wild-type JE2 grown in CDMG with sub-inhibitory OX (0.05 μg/ml), and not in the *pgl* mutant.

Not unexpectedly, the *pgl* mutation significantly perturbed the metabolome. However, accumulation of several individual metabolites within the PPP and glycolysis was not uniformly affected suggesting that the restoration of homeostasis required to enable growth in the absence of Pgl was complex. For example, downstream of Pgl in the PPP, levels of ribose-5-P were increased whereas sedoheptulose 7-P and nucleotide levels were decreased. The accumulation of ribose-5-P in a mutant lacking the transketolase *tkt* gene from the non-oxidative phase of the PPP was also accompanied by decreased sedoheptulose 7-P, although levels of inosine-5-monophosphate, xanthosine-5-monophosphate, and hypoxanthine were increased in the *tkt* mutant (48). The reduction in purine (and pyrimidine) nucleotide accumulation in the *pgl* mutant is consistent with its sensitivity to sulfamethoxazole, and with previous studies showing that mutations in the purine biosynthesis and salvage pathways are accompanied by increased OX resistance (26, 53). The metabolomics data presented here suggest that mutation of *pgl* was accompanied by a complex and intricately regulated interconversion of glycolytic and PPP intermediates to ensure maintenance of key central metabolites needed to support growth.

Analysis of PG architecture, crosslinking and overall concentration revealed no differences between the wild type and *pgl* strains suggesting that other changes in the cell envelope are responsible for increased *pgl* OX resistance. In this context reduced WGA binding to *pgl* cells, which is indicative of reduced teichoic acid levels or altered WTA/LTA structure, is of particular interest. Consistent with this *pgl* resistance to Congo red, which targets lipoteichoic acid synthase, LtaS, was also reduced. On the one hand, these observations correlate with the altered susceptibility of *pgl* to antimicrobial agents targeting WTAs and LTAs, and the ruffled morphology of the cell surface imaged by TEM. On the other hand, it is unclear how reduced levels of teichoic acids might be associated with increased β-lactam resistance. One possibility is that WGA binding to wild-type JE2 cells in CDMG OX may be unpredictably influenced by the extensive cell lysis, which is not observed in *pgl* mutant cells, which are smaller and have thick cell walls and intact septa. A second possibility is that WTA or LTA polymer length and/or post-translational modifications are altered in the *pgl* mutant. While construction of *pgl*/*tarM* and *pgl*/*tarS* double mutants excluded a role for α and β glycosylation of WTAs, respectively, we cannot rule out a possible role(s) for WTA/LTA polymer length, stability or release in the *pgl* OX resistance phenotype. Strikingly, mutations in VraFG/GraSR reversed the increased OX resistance phenotype of *pgl* in CDMG, as well as increased VAN resistance in MHB and reduced Congo red resistance. Meehl *et al* previously proposed that because mutation of *vraG* increased susceptibility to the structurally dissimilar vancomycin and polymyxin B in *S. aureus* strains Mu50 and COL, VraFG may play a broader role in the export of cell wall/teichoic acid precursors or modifying subunits, rather than specific antimicrobial agents (71). D-alanylation of WTAs was also shown to be reduced in a *graRS* mutant (72), further implicating this multienzyme membrane complex in cell envelope biogenesis. Importantly the restoration of wild-type OX MICs in CDMG in *pgl*/*vraF*, *pgl*/*vraG* and *pgl*/*graR* mutants was also accompanied by the restoration of WGA cell binding to wild-type levels, supporting a role for WTAs/LTAs in the increased OX resistance phenotype of the *pgl* mutant.

GraSR was also shown to regulate the transcription of *mprF*, which codes for the LysPG flippase implicated in resistance to CAMPs and daptomycin (79–81). Notably, a *mprF* missense mutation associated with increased cell size and daptomycin resistance was also shown to reduce MRSA OX resistance (82) raising the possibility that altered MprF activity could contribute to *pgl* phenotypes in a VraFG/GraRS-dependent manner.

The GraRS/VraFG complex shares significant amino acid sequence similarity with BceRS/BceAB in *Bacillus subtilis,* which plays an important role in bacitracin resistance. Bacitracin targets the lipid II cycle intermediate undecaprenyl-pyrophosphate (UPP), which is believed to be flipped/transported across the membrane, potentially by the BecAB ABC transporter, during PG biosynthesis (83, 84). Upregulation of *bceAB* expression by the BceR response regulator and changes in the conformation of BceAB appear to protect UPP from bacitracin inhibition (83). Thus, while PG structure and crosslinking is unaffected by the *pgl* mutant, it is tempting to speculate that UPP flipping across the membrane by VraFG may be of particular importance for PG biosynthesis in the *pgl* mutant under OX stress in CDMG, which is detected by the GraRS two component system. GraRS is known to be required for *S. aureus* growth at high temperatures and under oxidative stress (85), supporting the conclusion that the *vraFG*-dependent increase in OX resistance in *pgl* is also environmentally-regulated, as evidenced by changes in OX MICs in different culture media.

Taken together, our data reveal dramatically increased OX susceptibility and lysis of wild-type JE2 in CDMG, which is apparently due to the fragility of its cell envelope under these growth conditions. This vulnerability is reversed by the *pgl* mutation and the increased OX MIC and resistance to OX-induced lysis of the *pgl* mutant is, in turn, dependent on VraFG and GraRS. Mechanistically, the phenotypic consequences of metabolic reprogramming in the *pgl* mutant include increased flux to cell envelope precursors, altered susceptibility to drugs targeting WTAs and LTAs, reduced levels of teichoic acids, and cells that are smaller with a ruffled surface morphology thick cell walls and intact septa. These phenotypic changes in the cell envelope are apparently dependent on the VraFG/GraRS complex and we propose a possible model (Fig. 9), in which multienzyme membrane complex may directly and/or indirectly control the activity of the PG, WTA and LTA biosynthetic machinery, particularly in CDMG, to increase β-lactam resistance and prevent the extensive OX-induced lysis evident in the wild-type JE2.

**Fig. 9.**
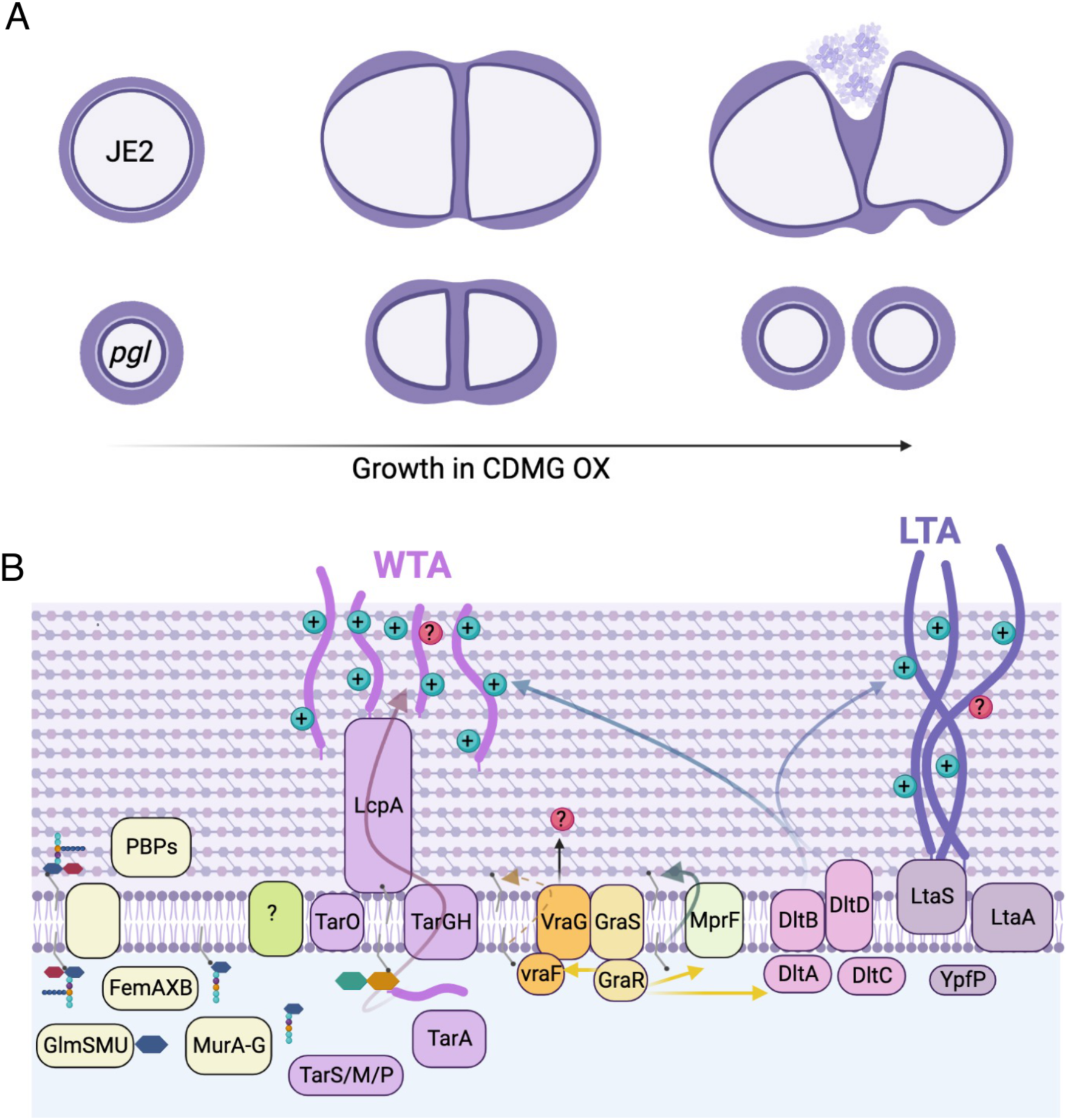
Suggested model for VraFG-dependent high-levelβ-lactam resistance in the MRSA *pgl* mutant. **A.** Illustration of JE2 and *pgl* cell division during growth in CDMG OX. *pgl* cells are smaller than wild-type JE2 when grown in CDMG and undergo normal cell division, whereas extensive lysis is evident among wild type cells. **B.** Illustration of peptidoglycan (PG), wall teichoic acid (WTA) and lipoteichoic acid (LTA) biosynthesis in a *pgl* mutant. Mutations in *vraF, vraG* and to a lesser extent *graR* reverse the increased OX resistance phenotype of the *pgl* mutant. Metabolic reprogramming in the *pgl* mutant increases carbon flux to cell envelope precursors and β-lactam resistance via a mechanism dependent on VraFG/GraRS-controlled regulation of WTA/LTA biosynthesis, export or posttranslational modification. Previous studies have implicated the VraFG/GraRS complex in resistance to cationic antimicrobial peptides and regulation of *dltABCD* and *mprF* transcription, and it has also been proposed to play a role in the export of peptidoglycan or teichoic acid precursors or modifying subunits.

## Materials and Methods

### Bacterial strains and growth conditions

Bacterial strains and plasmids used in this study can be found in Table S1. *Escherichia coli* strains were grown in Luria Bertani (LB) broth or agar (LBA) and *Staphylococcus aureus* strains were grown in Tryptic Soy Broth (TSB), Tryptic Soy Agar (TSA), Mueller-Hinton Broth (MHB) supplemented with 2% NaCl where indicated, Mueller-Hinton Agar (MHA) supplemented with 2% NaCl where indicated, Brain Heart Infusion (BHI) broth, LB broth, chemically defined medium (CDM), chemically defined medium supplemented with glucose (5 g/L) (CDMG). Culture media were supplemented with erythromycin (Erm) 10 μg/ml, chloramphenicol (Cm) 10 μg/ml, ampicillin (Amp) 100 μg/ml, kanamycin (Km) 90 μg/ml, oxacillin (OX) at varying concentrations as indicated.

### Genetic manipulation of *S. aureus,* complementation of NE202 (*pgl*) and construction of *pgl* double and triple mutants

To verify the increased OX resistance phenotype of NE202, phage 80α was used to transduce the *pgl*::Erm^r^ allele into wild-type JE2, as described previously (39, 57). The presence of the *pgl*::Erm^r^ allele in NE202 and the transductant was verified by PCR amplification with primers NE202_check_F, NE202_check_R, Martn_ermF, and Martn_ermR (Table S2).

To complement NE202, the *pgl* gene including its promoter and upstream regulatory sequences was first amplified from JE2 genomic DNA using *pgl*_Fwd and *pgl*_Rev primers (Table S2), cloned into pDrive (Qiagen) in *E. coli* TOP10 (Invitrogen), verified by Sanger sequencing (Source Biosciences) before being sub-cloned on an *Eco*RI restriction fragment into the *E. coli*-*Staphylococcus* shuttle plasmid pLI50 (86) and transformed into *E. coli* HST08 (Takara Bio). The pLI50_*pgl* plasmid was then isolated and transformed by electroporation into the restriction-deficient strain RN4220, and subsequently into NE202. All plasmid-harbouring strains were cultured in media supplemented with 100 μg/ml ampicillin (*E. coli*) or 10 μg/ml chloramphenicol (*S. aureus*) to maintain plasmid selection.

To generate the *pgl*::Km^r^ mutant, pKAN plasmid (57) was isolated from IM08B and electroporated into NE202 (*pgl*::Erm^r^) and the Erm^r^ marker swapped for the Km^r^ marker using allelic exchange (57). To construct a markerless Δ*pgl* mutant, the pTNT plasmid (57) from RN4220 pTNT was isolated and electroporated into NE202 (*pgl*::Erm^r^) and the Erm^r^ marker swapped out for a shorter, markerless version of the transposon insertion leaving a small deletion in the *pgl* gene. The *pgl*::Km^r^ and Δ*pgl* mutants were verified by PCR using primers NE202_check_F, NE202_check_R, KanR_fwd and KanR_rev (Table S2.)

To construct *pgl* double mutants phage 80α used to transduce the Erm^r^-marked alleles from the following Nebraska transposon library (57) mutants into *pgl*::Km^r^: NE1868 (*mecA*), NE952 (*gntP*), NE1124 (*gntK*), NE569 (*sucC*), NE547 (*sucA*), NE76 (*leuB*), NE239 (*putA*), NE1518 (*gudB*), NE70 (*vraG*), NE645 (*vraF*), NE481 (*graR*), NE942 (*tarS*), NE611 (*tarM*) NE626 (*sdhA*). The presence of transposon insertions in the genes was confirmed by PCR using primers listed in Table S2.

To construct the *pgl/vraG/putA* triple mutant the NE239 *putA*::Em^r^ strain was first transformed with pSPC plasmid (57) isolated from IM08B pSPC and allelic exchange performed as previously described (57) to generate *putA*::Spec^r^. The *putA*::Spec^r^ allele was then transduced into the *pgl/vraG* double mutant using phage 80α. The presence of transposon insertions in *pgl*, *vraG* and *putA* genes were confirmed by PCR using primers listed in Table S2.

### Tecan growth curves

A Tecan Sunrise microplate instrument equipped with Magellan software was used to record data from growth experiments performed in 96-well plates. Cultures were streaked on TSA plates supplemented with antibiotics where needed and grown at 37°C for 24 hours. The next day, colonies were resuspended in 1 ml of PBS, before being washed in PBS. The PBS washed cell suspensions were adjusted to an OD_600_ of 0.2 in 1 ml of PBS and 10 μl inoculated into wells containing 190 μl growth media (MHB, LB, TSB, BHI, CDM, CDMG, CDM 10 μg/ml OX, CDMG 10 μg/ml OX, CDMG supplemented with potassium D-Gluconate (5 g/L) (with or without OX 10 μg/ml) (starting OD_600_ = 0.01) and were then incubated at 35-37°C for 24 h with shaking and OD_600_ recorded every 15 min intervals. For H_2_O_2_ sensitivity assays (Figure S3), CDMG and CDMG containing 500 μM H_2_O_2_ were inoculated at a starting OD_600_ of 0.05. Three independent biological replicates were performed for each strain and the resulting data plotted using GraphPad Prism software V9.

### Antibiotic disc diffusion susceptibility assays

Disk diffusion susceptibility testing was performed in accordance with Clinical Laboratory Standards Institute (CLSI) guidelines (87) and as previously described (53) with the slight modifications. Briefly, overnight cultures were diluted into 5 ml fresh TSB and grown for 3 h at 37°C with shaking at 200 rpm. The 3 h grown cultures were then adjusted to *A*_600_ = 0.5 and 150 μl of this suspension was swabbed evenly 3 times across the surface of an MHA plate (4 mm agar thickness). Six mm blank discs (OXOID) were spotted with 20 μl antibiotics (cefoxitin 1.5 mg/ml stock). Once dried, the discs were applied onto the MHA plates spread with culture suspension before incubation for times specified by CLSI guidelines for stated antibiotics at 37°C. Three independent measurements were performed for each strain and zone of inhibition was measured and recorded.

### Antibiotic minimum inhibitory concentration (MIC) measurements and synergy/checkerboard assays

MIC measurements by broth microdilutions were performed in accordance with CLSI methods for dilution susceptibility testing of staphylococci (88) with modifications. Briefly, strains were first grown at 37°C on MHA 2% NaCl for 24 h and 5 - 10 colonies were resuspended in 0.85% saline before being adjusted to 0.5 McFarland standard (*A*_600_ = 0.1). The cell suspension was then diluted 1:20 in PBS and 10 μl used to inoculate 100 μl media (MHB 2% NaCl / CDM / CDMG) containing serially diluted antibiotics (oxacillin, fosfomycin, targocil, tunicamycin, Congo red, amsacrine, DCS, vancomycin and sulfamethoxazole) in 96-well plates. The plates were incubated at 35°C for 24 h and MIC values were recorded as the lowest antibiotic concentration where no growth was observed. Checkerboard/synergy assays were performed as previously described, using (0-128 μg/ml) fosfomycin and (0-256 μg/ml) oxacillin as indicated on Fig S7.

### Genomic DNA (gDNA) extraction and Whole Genome Sequencing (WGS)

Genomic DNA (gDNA) extractions were performed using the Wizard® Genomic DNA Purification Kit (Promega) following pre-treatment of *S. aureus* cells with 10 μg/ml lysostaphin (Ambi Products LLC) at 37°C for 30 min. The genome sequencing for NE202 (*pgl*) was performed by MicrobesNG using an Illumina HiSeq platform and a 250-bp paired end read kit. DNA libraries for *pgl*::Km^r^ and *pgl*R1 were prepared using an Illumina Nextera XT DNA Library Prep kit, validating size distribution by gel electrophoresis, and bead-normalizing the libraries. An Illumina MiSeq v2 600 cycle kit was used for genome sequencing, generating 300-bp paired end reads. PhiX was used as a sequencer loading control. The CLC Genomics Workbench software (Qiagen Version 20) was used for genome sequencing analysis of the different strains, as described previously (89). As a reference genome, a contig was produced for wild-type JE2 by mapping Illumina reads onto the closely related USA300 FPR3757 genome sequence (RefSeq accession number NC_007793.1). The Illumina short read sequences from NTML mutants (57) of interest were then mapped onto the assembled JE2 sequence, and the presence of the transposon insertion confirmed. Single Nucleotide Polymorphisms (SNPs), deletions or insertions were identified where present.

### PBP2a Western blot analysis

PBP2a Western blots were performed as previously described (90) with slight modifications. Briefly, single colonies from wild-type JE2, *pgl* and *pgl*_comp_, MSSA strain 8325-4 (negative control) and HoR MRSA strain BH1CC (positive control) were inoculated in TSB overnight and grown at 37°C with 200 rpm shaking. The next day, day cultures were started at OD_600_ 0.05 in 50 ml TSB supplemented with 0.5 μg/ml OX except for 8325-4 which was grown with no OX supplementation, and BHICC with 50 or 75 μg/ml OX, and grown for 6 hours, with shaking (200 rpm). Samples were pelleted and resuspended in PBS to an *A*_600_ = 10. Six μl of lysostaphin (10 μg/ml) and 1 μl of DNase (10 μg/ml) was added to 500 μl of this concentrated cell suspension before being incubated at 37°C for 40 min. Next, 50 μl of 10% SDS was added, and the incubation continued for a further 20 min. The lysed cells were then pelleted in a microcentrifuge for 15 min, following which the protein-containing supernatant was collected and total protein concentration determined using the Pierce BCA Protein Assay Kit. For each sample, 8 μg total protein was run on a 7.5% Tris-Glycine gel, transferred to a PVDF membrane, and probed with anti-PBP2a (1:1000), followed by HRP-conjugated protein G (1:2000) and colorimetric detection with Opti-4CN Substrate kit. Three independent experiments were performed, and a representative image is shown.

### Peptidoglycan (PG) analysis

Wild-type JE2, *pgl* and *pgl*_comp_ were grown in MHB and MHB supplemented with oxacillin 0.5 μg/ml, CDMG and CDMG supplemented with OX 0.05 μg/ml. For each strain and growth condition tested, independent quadruplicate 50 ml cultures were grown in flasks at 37°C with 200 rpm shaking overnight and cell pellets were collected at 4°C at 7000 rpm. The pellets were then resuspended in PBS, pelleted at 10000 rpm and snap frozen in liquid nitrogen in 1.5 ml tubes. Peptidoglycan (PG) was extracted from wild-type JE2, *pgl* and *pgl*_comp_ from boiled samples as described previously (91). Once boiled, cell wall material was pelleted by ultracentrifugation and washed with water. Clean sacculi were digested with muramidase (100 μg/ml) and soluble muropeptides reduced using 0.5 M sodium borate pH 9.5 and 10 mg/mL sodium borohydride. The pH of the samples was then adjusted to 3.5 with phosphoric acid. UPLC analyses were performed on a Waters-UPLC system equipped with an ACQUITY UPLC BEH C18 Column, 130Å, 1.7 μm, 2.1 mm × 150 mm (Waters Corporation, USA) and identified at Abs. 204 nm. Muropeptides were separated using a linear gradient from buffer A (0.1% formic acid in water) to buffer B (0.1% formic acid in acetonitrile). Identification of individual peaks was assigned by comparison of the retention times and profiles to validated chromatograms. The relative amount of each muropeptide was calculated by dividing the peak area of a muropeptide by the total area of the chromatogram. The abundance of PG (total PG) was assessed by normalizing the total area of the chromatogram to the OD600. The degree of cross-linking was calculated as described previously (92).The data for at least three independent experiments were plotted using GraphPad Prism software.

### Confocal microscopy and cell size determination

For microscopy experiments, JE2, *pgl* and *pgl*_comp_ were grown overnight at 37°C in CDMG w/wo 0.05 μg/ml OX. The next day, the cultures were washed and normalized to an *A*_600_ of 1 in PBS and 75 μl of these cultures were double stained for 30 mins at 37°C with vancomycin-BODIPY FL at a final concentration of 2 μg/ml and WGA Alexa Fluor 594 at a final concentration of 25 μg/ml. Bacteria were then collected by centrifugation for 2 mins at 14,000 x*g*. The cells were resuspended with 100 μl of PBS, pH 7.4, and 5 μl of this sample was spotted onto a thin 1% agarose gel patch prepared in PBS. Stained bacteria were then imaged at X1000 magnification using Olympus LS FLUOVIEW Fv3000 Confocal Laser Scanning Microscope. Cell size was measured as previously described (54) using ImageJ software (Fiji v.1.0). Images of cells from three biological replicates were acquired, 50 cells measured per biological replicate (150-200 cells in total per condition), and the average and standard deviations for the three/four biological replicates were plotted using GraphPad Prism version 9.2 and significant differences were determined using a Kruskal-Wallis test followed by a Dunn’s multiple comparison test. Only 60 cells could be measured for OX treated cells due to cell lysis.

### Transmission Electron Microscopy (TEM) and cell morphology analysis

Overnight cultures of JE2, *pgl* and *pgl*_comp_ were grown overnight in 5 ml CDMG at 37°C shaking at 200 rpm. The next day, OD_600_ values were measured, and cultures were used to inoculate 5 ml day cultures in CDMG 1 μg/ml OX to OD_600_ of 0.06. The day cultures were grown for 4.5 hours at 35°C shaking at 200 rpm, before being pelleted down, and normalised to OD_600_ of 1 in PBS. Cells pellets were resuspended in 0.2M sodium cacodylate buffer pH 7.2. Fixed bacteria were dehydrated, embedded in resin, and thin sectioned in the University of Galway Centre for Microscopy & Imaging. Images were acquired using Hitachi H7500 Transmission Electron Microscope. Representative cells from each strain were imaged at 30,000× and 100,000× magnification.

### Congo Red susceptibility spotting assays

*S. aureus* strains JE2, *pgl*, *pgl*_comp_, *vraG* and *pgl/vraG* were streaked onto TSA plates containing appropriate antibiotics, and the plates were incubated overnight at 37°C. The next day, overnight cultures of the strains from single colonies were grown in 5 ml TSB, at 37°C shaking at 200 rpm. The next day, PBS washed cells were normalised to an OD_600_ of 1 per ml in PBS and serial dilutions prepared from 10^-1^ until 10^-8^ in a 96-well plate. Five μl of the serially diluted cell suspensions was spotted onto TSA plates containing 0.125% Congo Red. The plates were dried in a flow hood and were incubated at 37°C for 24 hours. Plates were visualised and photos were taken for three biological replicates. Representative image is shown in Figure 8B.

### Quantification of Wheat Germ Agglutinin (WGA) binding

Overnight cultures of *S. aureus* strains were grown in 3 ml CDMG at 37°C shaking at 200 rpm. The next day, OD_600_ values were measured, and cultures were used to inoculate 5 ml day cultures in CDMG 0.1 μg/ml OX to OD_600_ of 0.06. The day cultures were grown for 4.5 hours at 35°C shaking at 200 rpm, before being pelleted down, washed with PBS, and normalised to OD_600_ of 1 in PBS. One hundred μl of this cell suspension was incubated with WGA Alexa Fluor 594 at a final concentration of 25 μg/ml for 30 minutes at 37°C. After the incubation with the dye, the cells were pelleted at 14,000 rpm for 3 minutes, and the supernatant was used for fluorescence measurements in Polarstar plate reader (Excitation/Emission 590/617 nm). PBS containing WGA Alexa Fluor 594 at a final concentration of 25 μg/ml was used as a positive control, and PBS was used as a blank control. The reduction in WGA Alexa Fluor 594 from the positive control was calculated per sample, and % bound WGA plotted using were plotted and significant differences were determined for two biological repeats using two-way ANOVA with Turkey’s post-hoc. using GraphPad Prism version 9.2

### Culture supernatant sample preparation for LC-MS/MS

Overnight cultures of *S. aureus* strains were grown in 3 ml CDMG at 37°C shaking at 250 rpm. The next day, 250 ml flasks containing 25 ml CDMG were inoculated to an OD_600_ of 0.06 and were grown for 7.5 h (OD_600_= 4.22-4.96). One ml from the cultures were collected and centrifuged at 12,000 rpm, 10 min at 4°C, and supernatant collected. These samples were diluted 1:100 v/v using 10 mM NH_4_OAc + 10mM NH_4_OH + 5% acetonitrile. 5 μl was injected into the LC-MS/MS (details below).

### Sample preparation intracellular metabolite analysis by LC-MS/MS

Overnight cultures of *S. aureus* strains were grown in 3 ml CDMG at 37°C shaking at 250 rpm. The next day, 250 ml flasks containing 25 ml CDMG (with or without 1 μg/ml OX) were inoculated at a starting OD_600_ of 0.06 and grown for 4-5 hours (until exponential phase was reached) at 37°C shaking at 250 rpm. Culture volumes corresponding to OD_600_ of 10 were then harvested and rapidly filtered through a membrane (0.45 μm, Millipore). The cells on the membrane were washed twice with 5 ml cold saline and immediately quenched in ice-cold 60% ethanol containing 2 μM Br-ATP as an internal control. The cells were mechanically disrupted using a bead homogenizer set to oscillate for 3 cycles (30 s) of 6800 rpm with a 10 s pause between each cycle. Cell debris was separated by centrifugation at 12,000 rpm. The supernatant containing intracellular metabolites were lyophilized and stored at −80°C. These samples were reconstituted in 100 μl of 50% MeOH.

### Analysis of PPP flux using 1,2-^13^C glucose

The *S. aureus* strains were inoculated in 25 ml CDM containing either unlabelled or 1,2-^13^C-labeled glucose at a starting OD_600_ of 0.06. The cultures were grown at 37°C with shaking at 250 RPM until the OD_600_ reached 1.0. The culture volume corresponding to an OD_600_ of 10 was then harvested and immediately filtered through a 0.45 μm Millipore membrane before being subjected to further processing as outlined in the previous section.

### LC-MS/MS mass spectrometry

A triple-quadrupole-ion trap hybrid mass spectrometer (QTRAP6500+ by Sciex, USA) connected to an ultra-performance liquid chromatography I-class (UPLC) system (Waters, USA) was utilized for metabolomics analysis. The chromatographic separation was performed using the UPLC on a XBridge Amide analytical column (150 mm x 2.1 mm ID, 3.5 μm particle size by Waters, USA) and a binary solvent system with a flow rate of 0.3 ml/min. The analytical column was preceded by a guard XBridge Amide column (20 mm x 2.1 mm ID, 3.5 μm particle size by Waters, USA). The mobile phase A consisted of 10 mM ammonium acetate and 10 mM ammonium hydroxide with 5% acetonitrile in LC-MS grade water (pH adjusted to 8.0 with glacial acetic acid), while mobile phase B was 100% LC-MS grade acetonitrile. The column was maintained at 40°C and the autosampler temperature was kept at 5°C throughout the sample run. The gradient conditions were as follows: A/B ratio of 15/85 for 0.1 minute, 16/84 for 3.0 minutes, 35/65 for 4.0 minutes, 40/60 for 5.0 minutes, 45/55 for 3.0 minutes, 50/50 for 5.5 minutes, 30/70 for 1.5 minutes, and finally equilibrated at 15/85 for 5.0 minutes before the next run. The needle was washed with 1000 μL of strong wash solvent (100% acetonitrile) and 1000 μL of weak wash solvent (10% aqueous methanol) prior to injection, with an injection volume of 5 μL. The QTRAP6500+ operated in polarity switching mode for the targeted quantitation of amino acids through the Multiple Reaction Monitoring (MRM) process. The electrospray ionization (ESI) parameters were optimized, with an electrospray ion voltage of −4200 V in negative mode and 5500V in positive mode, a source temperature of 400°C, a curtain gas of 35 and gas 1 and 2 of 40 and 40 psi, respectively. Compound-specific parameters were optimized for each compound through manual tuning, with declustering potential (DP) of 65V in positive mode and - 60V in negative mode, entrance potential (EP) of 10V in positive mode and −10V in negative mode, and collision cell exit potential (CXP) of 10V in positive mode and - 10V in negative mode.

## Data availability

Whole-genome sequence data is available from the European Nucleotide Archive Registered Project PRJEB59981, sample accession numbers ERS14733509-ERS14733512 and run accession numbers ERR10960504-ERR10960507. The SAUSA300_FRP3757 (TaxID: 451515) reference genome sequence is available from NCBI.

## Acknowledgements

M.S.Z., L.A.G., A.N., O.B. and J.P.O’G. are supported by grants from the Health Research Board (ILP-POR-2019-102 and HRA-POR-2015-1158), Science Foundation Ireland (19/FFP/6441) and the Irish Research Council (GOIPG/2019/2011). F. R., M.S., J.A., D.S. P.D.T and V.C.T are financially supported by research grants P01 AI083211 (to P.D.T. and V.C.T) and R01 AI125588 (to V.C.T) from the National Institute of Allergy and Infectious Diseases, USA. E.B. and F.C. are supported by the Swedish Research Council, the Laboratory for Molecular Infection Medicine Sweden (MIMS), Umeå University, the Knut and Alice Wallenberg Foundation (KAW) and the Kempe Foundation. We are grateful to Dr Peter Owens and Dr Emma McDermott from the University of Galway Centre for Microscopy & Imaging (https://imaging.universityofgalway.ie) for their technical and scientific assistance with confocal and electron microscopy analysis, and to Volkhard Kaever, Hannover Medical School for preliminary metabolomic analysis.

The funders had no role in study design, data collection and interpretation, or the decision to submit the work for publication.

M.S.Z. and J.P.O’G. conceptualized the study. Formal analysis was performed by M.S.Z., L.A.G., E.B., A.C.N., J.A., E.O’N., F.R., P.D.F., F.C., V.C.T and J.P.O’G. The investigation and methodology was performed by M.S.Z., L.A.G., E.B., A.C.N., J.A., D.S., M.S., Ó.B. and F.R. The data was curated by M.S.Z. The original draft of the manuscript was written by M.S.Z. and J.P.O’G. and was reviewed, edited and approved by all authors. Funding was acquired by E.O’N., P.D.F., F.C., V.C.T. and J.P.O’G. The project was administered by J.P.O’G.

