## Supplemental Tables and Figures for "Metabolic reprogramming and flux to cell envelope precursors in a pentose phosphate pathway mutant increases MRSA resistance to β-lactam antibiotics"

**Table S1.** Bacterial strains and plasmids used in this study

| Strains/plasmids | Relevant Details |
|---|---|
| ***Staphylococcus aureus* strains** | |
| JE2 | USA300 cured of p01 & p03. Parent strain from the Nebraska Transposon Mutant Library (NTML) (57) |
| BH1CC | MRSA clinical isolate; SCC*mec* type II; CC8 (93) |
| 8325-4 | NCTC 8325 derivative cured of prophages (94), MSSA, CC8. |
| RN4220 | Restriction deficient derivative of *S. aureus* 8325 (94) |
| RN4220 pTNT | RN4220 carrying pTNT plasmid. $Cm^r$. (95) |
| NE202 (*pgl*) | JE2 *pgl*::$Erm^r$ (SAUSA300_1902). $Erm^r$. (57) |
| *pgl*::$Km^r$ | JE2 *pgl*::Km, $Km^r$ |
| *pgl*R1 | JE2 Δ*pgl*, VraG $Gln_{394}$STOP |
| NE202 pLI50_*pgl* (*pgl*$_{comp}$) | NE202 pLI50_*pgl* $Erm^r$, $Cm^r$ |
| NE952 (*gntP*) | JE2 *gntP*. $Erm^r$. (57) |
| NE1124 (*gntK*) | JE2 *gntK*. $Erm^r$. (57) |
| NE569 (*sucC*) | JE2 *sucC*. $Erm^r$. (57) |
| NE547 (*sucA*) | JE2 *sucA*. $Erm^r$. (57) |
| NE76 (*leuB*) | JE2 *leuB*. $Erm^r$. (57) |
| NE239 (*putA*) | JE2 *putA*. $Erm^r$. (57) |
| NE1518(*gudB*) | JE2 *gudB*. $Erm^r$. (57) |
| NE70 (*vraG*) | JE2 *vraG*. $Erm^r$. (57) |
| NE645 (*vraF*) | JE2 *vraF*. $Erm^r$. (57) |
| NE481 (*graR*) | JE2 *graR*. $Erm^r$. (57) |
| NE1868 (*mecA*) | JE2 *mecA*. $Erm^r$. (57) |
| NE626 *(sdhA)* | JE2 *sdhA*. $Erm^r$. (57) |
| NE942 (*tarS*) | JE2 *tarS*. $Erm^r$. (57) |
| NE611 (*tarM*) | JE2 *tarM*. $Erm^r$. (57) |
| JE2 *pgl*::$Erm^r$ | JE2 transductant. *pgl*::$Erm^r$. This study. |
| JE2 *gntP*::$Erm^r$ | JE2 transductant. *gntP*::$Erm^r$. This study. |
| JE2 *gntK*::$Erm^r$ | JE2 transductant. *gntK*::$Erm^r$. This study. |
| *pgl/gntP* | $Km^r$, $Erm^r$. This study. |
| *pgl/gntK* | $Km^r$, $Erm^r$. This study. |
| *pgl/sucC* | $Km^r$, $Erm^r$. This study. |
| *pgl/sucA* | $Km^r$, $Erm^r$. This study. |
| *pgl/leuB* | $Km^r$, $Erm^r$. This study. |
| *pgl/putA* | $Km^r$, $Erm^r$. This study. |
| *pgl/gudB* | $Km^r$, $Erm^r$. This study. |
| *pgl/vraG* | $Km^r$, $Erm^r$. This study. |
| *pgl/vraF* | $Km^r$, $Erm^r$. This study. |
| *pgl/graR* | $Km^r$, $Erm^r$. This study. |
| *pgl/thrC* | $Km^r$, $Erm^r$. This study. |
| *pgl/mecA* | $Km^r$, $Erm^r$. This study. |
| *pgl/sdhA* | $Km^r$, $Erm^r$. This study. |
| *pgl/putA*$_{Spec}$ | $Km^r$, $Spec^r$. This study. |
| *pgl/putA/vraG* | $Km^r$, $Spec^r$, $Erm^r$. This study. |
| *pgl/tarS* | $Km^r$, $Erm^r$. This study. |
| *pgl/tarM* | $Km^r$, $Erm^r$. This study. |
| ***Escherichia coli* strains** | |
| TOP10 | (F- *mcrA* Δ(*mrr-hsdRMS-mcrBC*) φ80*lacZ*ΔM15 Δ*lacX*74 *nupG recA1 araD139* Δ(araleu)7697 *galE15 galK16 rpsL*(StrR ) *endA1* λ (Invitrogen) |

| | |
|---|---|
| TOP10 pDrive | *E.coli* TOP10 carrying pDrive_*pgl*. This study. |
| HST08 | F- ,*endA1*, *supE44*,*thi*-1,*recA1*,*relA1*,*gyrA96*,*phoA*, Φ80d*lacZ*ΔM15, Δ (*lacZYA -argF* ) U169, Δ (*mrr -hsdRMS - mcrBC* ), *ΔmcrA*,λ– (Takara Bio) |
| HST08 pLI50_*pgl* | *E. coli* HST08 carrying pLI50_*pgl*. Amp$^r$. This study. |
| IM08B | SA08BΩPN25-*hsdS* (CC8-1) (SAUSA300_0406) of NRS384 integrated between the *essQ* and *cspB* genes (96) |
| IM08B pKAN | IM08B carrying pKAN. Amp$^r$. This study. |
| IM08B pSPC | IM08B carrying pSPC. Amp$^r$. This study. |
| **Plasmids** | |
| pDrive | *E. coli* cloning vector (Qiagen) |
| pDrive_*pgl* | pDrive carrying *pgl* from JE2. This study. |
| pLI50 | *E. coli* (Amp$^r$)-*Staphylococcus* (Cm$^r$)shuttle vector (86) |
| pLI50_*pgl* | pLI50 carrying *pgl* from JE2. *E. coli* (Amp$^r$)-*Staphylococcus* (Cm$^r$). This study. |
| pTNT | pJB38 with homologous DNA to *bursa aurealis* (95) |
| pKAN | pTNT with *aphA-3* (95) |
| pSPC | pTNT with *aad9* (95) |

**Table S2.** Oligonucleotide primers used in this study.

| Target gene | Primer name | Primer sequence (5'-3') |
|---|---|---|
| *pgl* | *pgl*_Fwd | TCATCCTTAATTCACCCCAATC |
| | *pgl*_Rev | CAGGTGTCCATTTACCACCA |
| | NE202_check_F | CCTAGGGTGCCGTCTCAGCCTTGGTCTTCG |
| | NE202_check_R | TCTGAGTTGACGCCTAATGTTGCACGAGTG |
| *gntP* | NE952_check_F | ACATCGATCATTACAGCGTTAATGCTA |
| *gntK* | NE1124_check_F | GAAGAAACAACTTGAAATGATGAAAGTG |
| *mecA* | NE1868_check_F | GGTGAAGTAGAAATGACTGAACGTC |
| $Erm^r$ | Martn_ermF | TTTATGGTACCATTTCATTTTCCTGCTTTTTC |
| | Martn_ermR | AAACTGATTTTTAGTAAACAGTTGACGATATTC |
| $Kan^r$ | KanR_fwd | GACCTAGGGGTTTCAAAATCGGCTC |
| | KanR_rev | GGCCTAGGTACTAAAACAATTCATCCAGTAAA |
| *graR* | NE481_check_F | GTTGCTGGTATTGAAGATTTCGG |
| *tarS* | NE942_check_F | CGATCAAGTGAGCGTTTAGTCAG |
| *tarM* | NE611_check_R | CAGCACCATTATTAGCATTAATATTCCTTG |
| *thrC* | NE886_check_R | GAATCGCTAAAATATCAGGTGCTTC |
| *gudB* | NE1518_check_R | CACCTAGTGCAGTTGATCTGTCG |
| *sdhC* | NE626_check_F | GCACATGTAGATTTGTTCTCAGTTGTACC |
| *leuB* | NE76_check_F | CTGTCACTGAAGGTACTGATGCCCAAGC |
| *putA* | NE239_check_R | GGTACTTATCAACTAATTCGTGGCTATCG |
| *vraG* | NE70_check_F | TGGTAACGCATGATCCTGTTGCAGCAAGC |
| *vraF* | NE645_check | CAGCTGATGTTCGTTGCCTTTGTCCACCAGAC |
| *sucC* | *sucC*_F | TACTCAAATCGCCATGCAGC |
| | *sucC*_R | AATGACTGAAACCGTTGCCC |
| *sucA* | *sucA*_F | GGCGGTAATGGACTCGGATT |
| | *sucA*_R | TCTACGCTATCCCCTACGTT |
| **Cloning primers** | | |
| *pgl* | *pgl*_F | TCATCCTTAATTCACCCCAATC |
| | *pgl*_R | CAGGTGTCCATTTACCACCA |

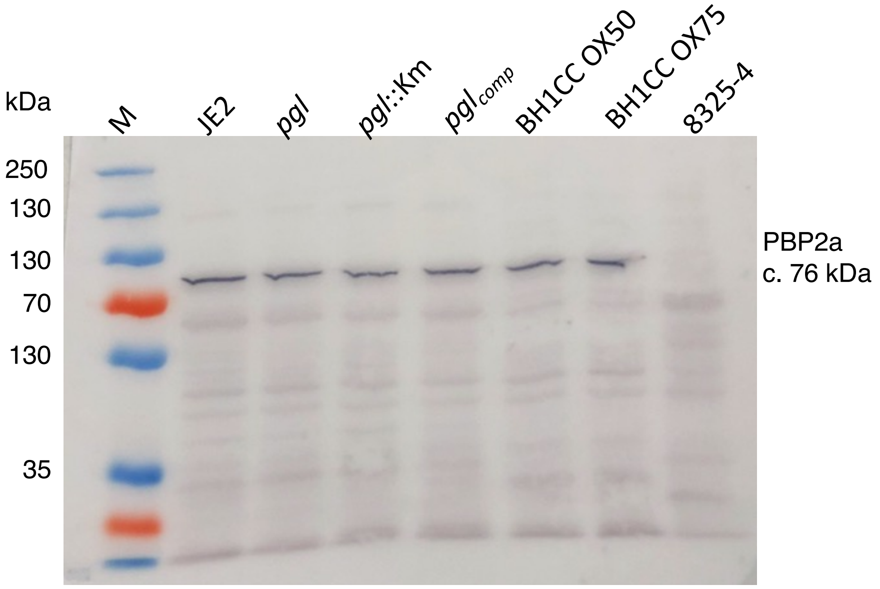


**Fig. S1.** Western blot of PBP2a protein in JE2, NE202 (*pgl*), *pgl*::$Km^r$, $pgl_{comp}$, HoR MRSA strain BH1CC (positive control) grown in OX 50, BH1CC grown in OX 75, MSSA strain 8325-4 (negative control). The JE2 and *pgl* strains were grown for 6 hours in TSB supplemented with 0.5 $\mu$g/ml oxacillin (OX), BHICC was grown in TSB OX 50 or 75 $\mu$g/ml, and 8325-4 which was grown in TSB with no OX. For each sample, 8 $\mu$g total protein was run on a 7.5% Tris-Glycine gel, transferred to a PVDF membrane and probed with anti-PBP2a (1:1000), followed by HRP-conjugated protein G (1:2000) and colorimetric detection with Opti-4CN Substrate kit. Three independent experiments were performed, and a representative blot is shown.

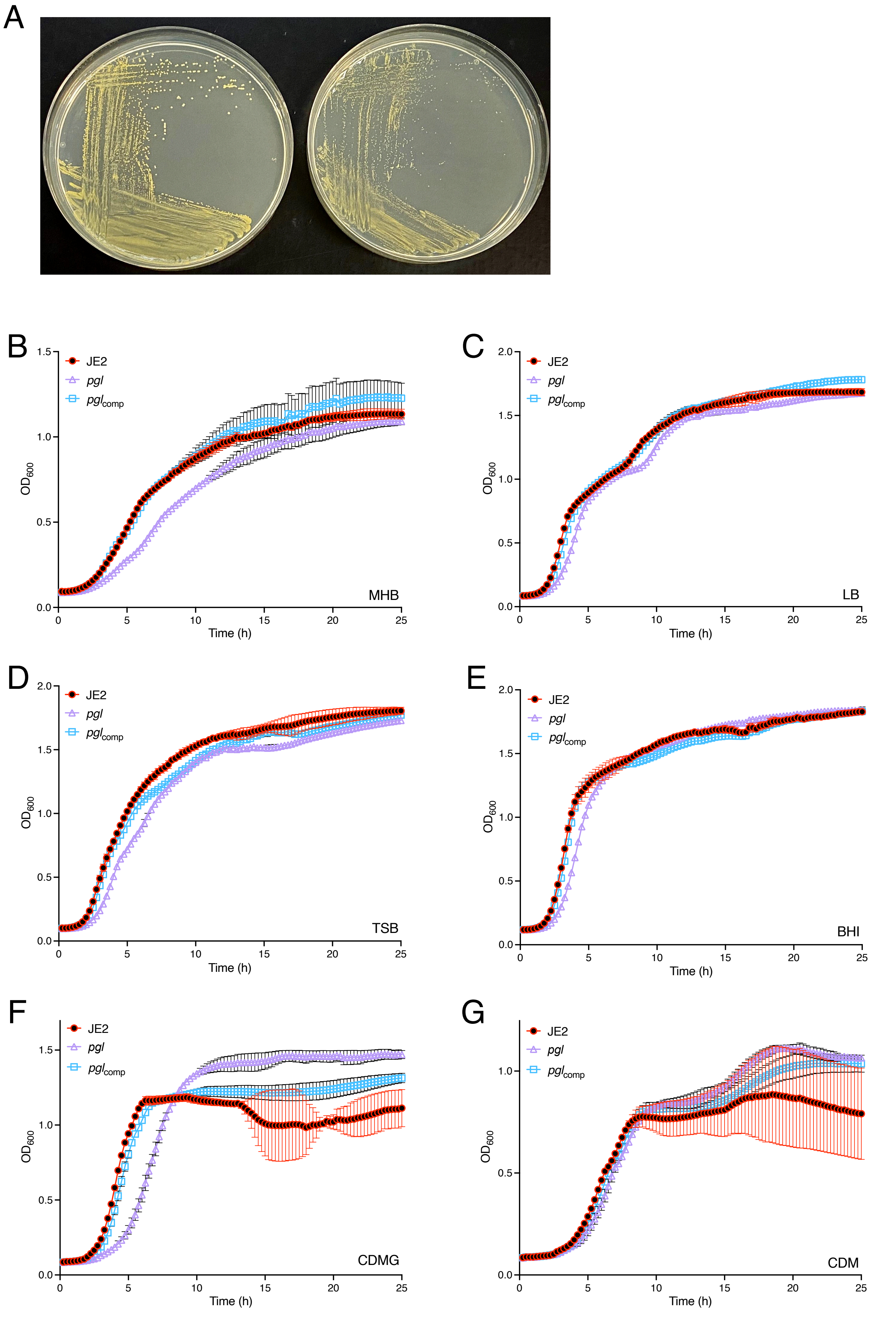

A
B
C
D
E
F
G
JE2
pgl
pgl comp
OD600
Time (h)
MHB
LB
TSB
BHI
CDMG
CDM

**Fig. S2. Impact of the *pgl* mutation on growth in different culture media. A.** Isolated colonies of JE2 (left) and *pgl* (right) after growth on MHA for 24 h at 37°C. **B-G.** Growth of JE2, *pgl* and the complemented *pgl* mutant for 25 hrs at 37°C in Mueller Hinton broth, MHB (B), Luria Bertani, LB (C), Tryptic Soya broth, TSB (D), Brain Heart Infusion, BHI (E), Chemically defined media with glucose, CDMG (F) and chemically defined media with no glucose, CDM (G). Growth ($OD_{600}$) was measured at 15 min intervals in a Tecan plate reader. Data are the average of 3 independent experiments plotted using GraphPad Prism V9 and error bars represent standard deviation.

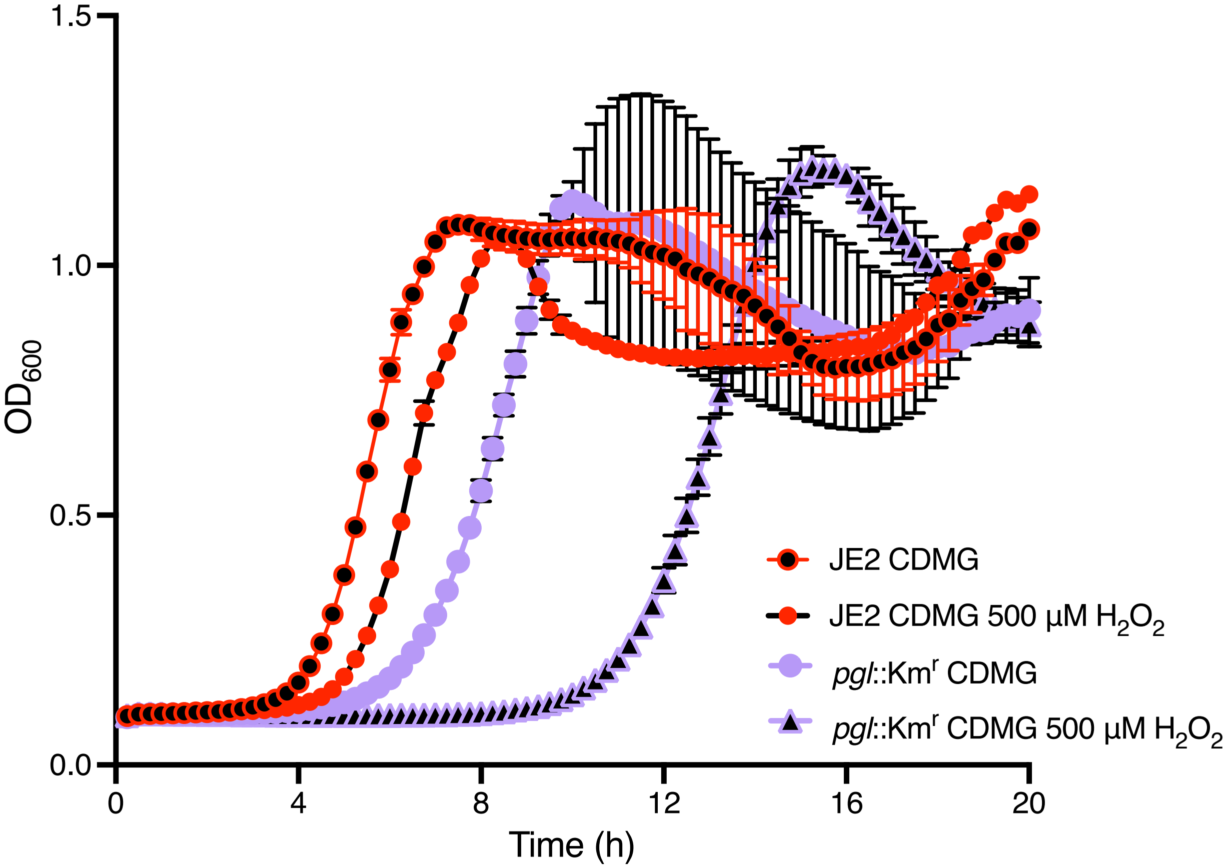


**Fig. S3. Mutation of *pgl* increases sensitivity to oxidative stress.** Growth of wild-type JE2 and *pgl*::Km$^r$ for 24 hrs at 37$^o$C in CDMG or CDMG supplemented with 500 µM $H_2O_2$. Growth ($OD_{600}$) was measured at 15 min intervals in a Tecan plate reader. Data are the average of 3 independent experiments plotted using GraphPad Prism V9 and error bars represent standard deviation.

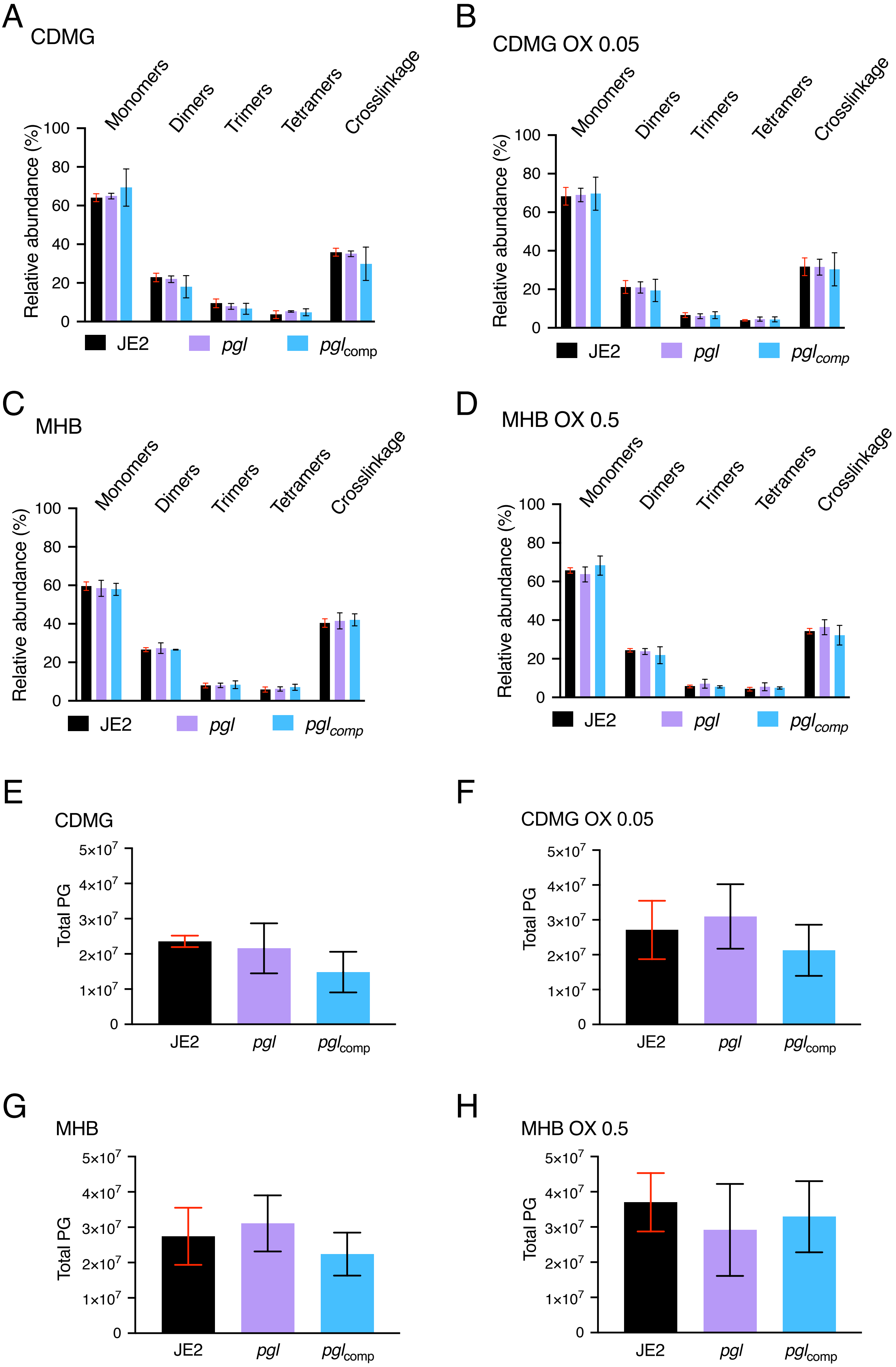
A
CDMG
B
CDMG OX 0.05
C
MHB
D
MHB OX 0.5
Monomers
Dimers
Trimers
Tetramers
Crosslinkage
Relative abundance (%)
0
20
40
60
80
100
JE2
pgl
pgl_comp
E
CDMG
F
CDMG OX 0.05
G
MHB
H
MHB OX 0.5
Total PG
0
1x10^7
2x10^7
3x10^7
4x10^7
5x10^7

**Fig. S4. Comparison of peptidoglycan oligomerisation, cross-linking and total amount in JE2, *pgl* and the complemented *pgl* mutant. A-D.** Relative proportions of cell wall muropeptide fractions based on oligomerization relative cross-linking efficiency in peptidoglycan extracted from JE2, *pgl* and $pgl_{comp}$ grown to exponential phase in CDMG (A), CDMG supplemented with OX 0.05 μg/ml (B), MHB (C) and MHB supplemented with OX 0.5 mg/ml (D). **E-H.** Total peptidoglycan (PG) extracted from normalised cell extracts of JE2, *pgl* and $pgl_{comp}$ grown to exponential phase in CDMG (E), CDMG supplemented with OX 0.05 μg/ml (F), MHB (G) and MHB supplemented with OX 0.5 μg/ml (H). The total PG content was calculated as the area below the chromatogram peaks/$OD_{600}$ and mean and standard deviation from three/four biological repeats plotted using GraphPad Prism V9.

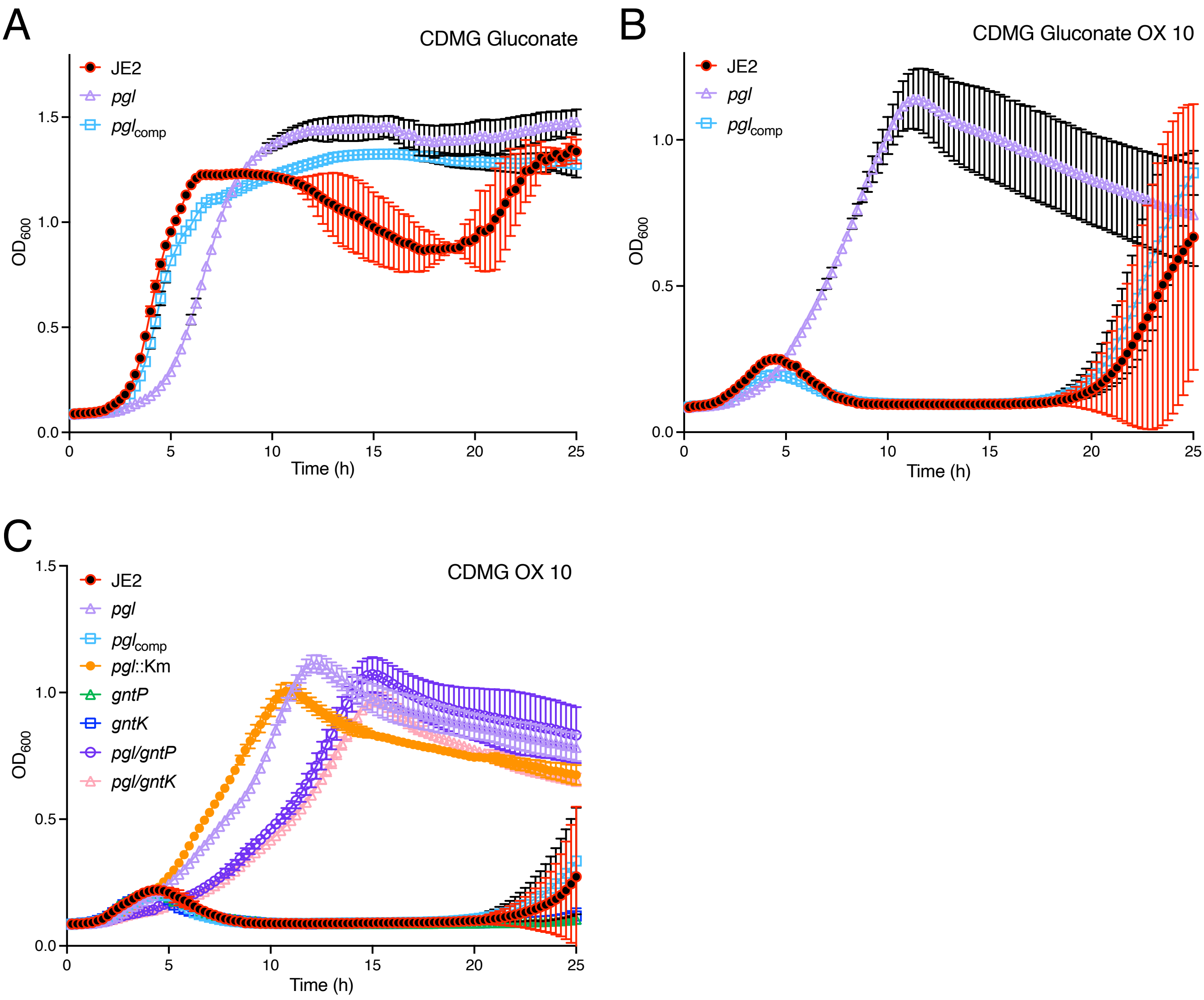


**Fig. S5. Exogenous addition of D-gluconate or mutation of the gluconate shunt genes *gntP* or *gntK* did not impact growth or the increased OX resistance of the *pgl* mutant. A and B.** Growth of JE2, *pgl* and the complemented *pgl* mutant for 25 hrs at 35ºC in CDMG supplemented with 5g/l potassium D-gluconate (0.5%) and no OX (A) or with both gluconate (0.5%) and OX 10 µg/ml (B). **C.** Growth of JE2, *pgl, pgl*$_{comp}$ *pgl*::Km$^r$, *gntP* (NE952), *gntK* (NE1124), *pgl/gntP* and *pgl/gntK* for 25 hrs at 35ºC in CDMG supplemented with OX 10 µg/ml. Growth ($OD_{600}$) was measured at 15 min intervals in a Tecan plate reader. Data are the average of 3 independent experiments using GraphPad Prism V9 and error bars represent standard deviation.

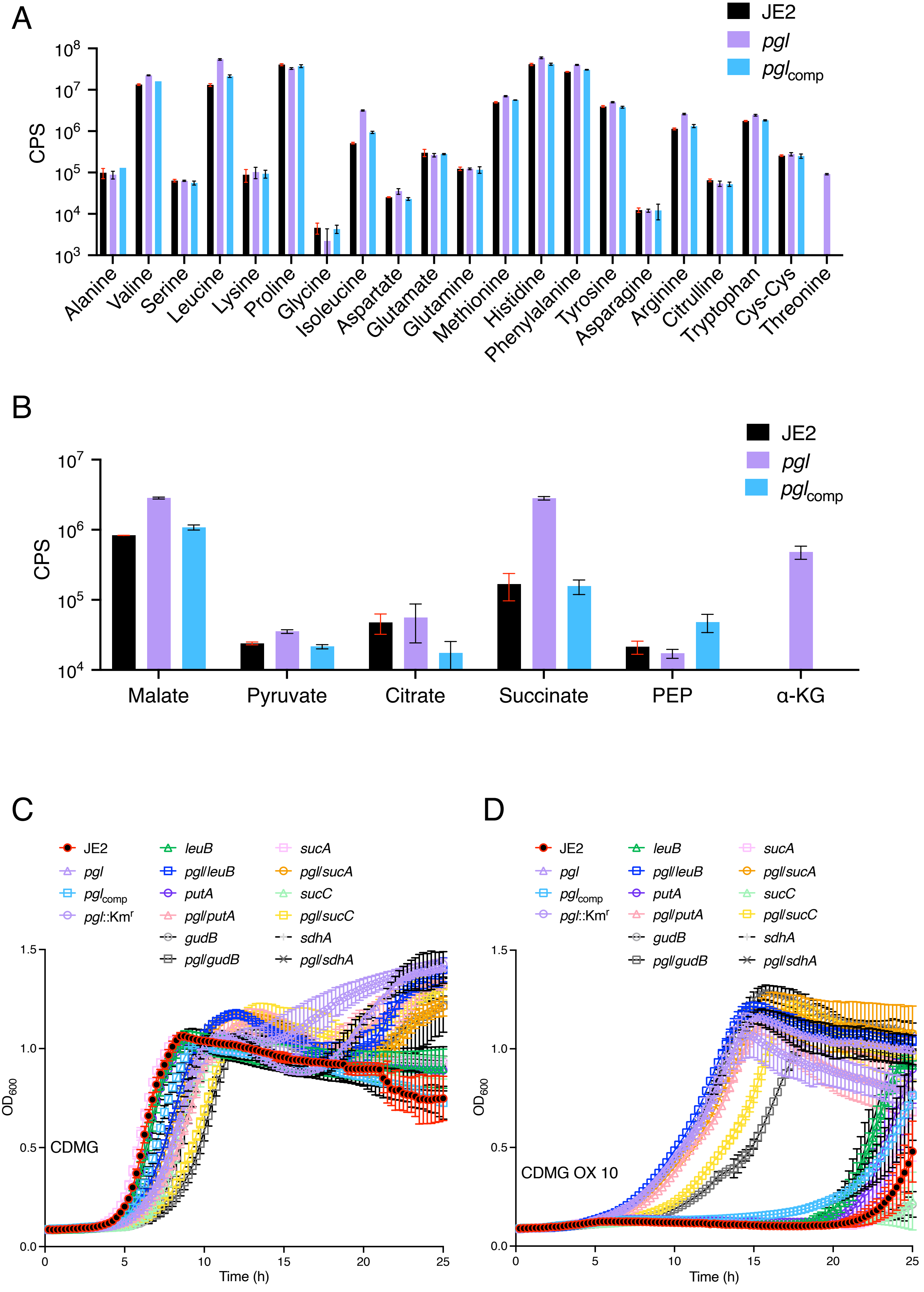

A
JE2
pgl
pglcomp
CPS
10^8
10^7
10^6
10^5
10^4
10^3
Alanine
Valine
Serine
Leucine
Lysine
Proline
Glycine
Isoleucine
Aspartate
Glutamate
Glutamine
Methionine
Histidine
Phenylalanine
Tyrosine
Asparagine
Arginine
Citrulline
Tryptophan
Cys-Cys
Threonine
B
JE2
pgl
pglcomp
CPS
10^7
10^6
10^5
10^4
Malate
Pyruvate
Citrate
Succinate
PEP
α-KG
C
JE2
pgl
pglcomp
pgl::Km^r
leuB
pgl/leuB
putA
pgl/putA
gudB
pgl/gudB
sucA
pgl/sucA
sucC
pgl/sucC
sdhA
pgl/sdhA
OD600
CDMG
Time (h)
D
JE2
pgl
pglcomp
pgl::Km^r
leuB
pgl/leuB
putA
pgl/putA
gudB
pgl/gudB
sucA
pgl/sucA
sucC
pgl/sucC
sdhA
pgl/sdhA
OD600
CDMG OX 10
Time (h)

**Fig. S6. Mutations in amino acid and TCA cycle biosynthetic genes did not reverse increased OX resistance in the *pgl* mutant**. **A and B.** Comparison of amino acid (A) and TCA cycle metabolites (B) in supernatants of JE2, *pgl* and $pgl_{comp}$ cultures grown for 7.5 h in CDMG measured by HPLC. Cell densities ($OD_{600}$) were normalized to each other before the cells were pelleted and supernatants collected. The data (CPS) shown are the average of three independent experiments and standard deviations are shown. **C and D.** Growth of JE2, *pgl, pgl*::Km$^r$, $pgl_{comp}$, *putA*, *gudB*, *leuB*, *sdhA*, *sucA*, *sucC*, *pgl*/*putA*, *pgl*/*gudB*, *pgl*/*leuB*, *pgl*/*sdhA*, *pgl*/*sucA* and *pgl*/*sucC* for 25 hrs at 35$^o$C in CDMG (C) and CDMG supplemented with OX 10 μg/ml (D). Growth ($OD_{600}$) was measured at 15 min intervals in a Tecan plate reader. Data are the average of 3 independent experiments using GraphPad Prism V9 and error bars represent standard deviation.

A

**JE2**

Oxacillin ($\mu$g/ml) →

| Fosfomycin ($\mu$g/ml) | 0 | 0.25 | 0.5 | 1 | 2 | 4 | 8 | 16 | 32 | 64 | 128 | 256 |
|---|---|---|---|---|---|---|---|---|---|---|---|---|
| 128 | 0.053 | 0.05 | 0.051 | 0.048 | 0.049 | 0.046 | 0.048 | 0.046 | 0.049 | 0.05 | 0.053 | 0.056 |
| 64 | 0.049 | 0.048 | 0.048 | 0.045 | 0.046 | 0.044 | 0.046 | 0.046 | 0.047 | 0.048 | 0.052 | 0.057 |
| 32 | 0.054 | 0.046 | 0.046 | 0.046 | 0.047 | 0.045 | 0.046 | 0.044 | 0.044 | 0.046 | 0.049 | 0.055 |
| 16 | 0.204 | 0.046 | 0.047 | 0.047 | 0.047 | 0.045 | 0.046 | 0.044 | 0.044 | 0.045 | 0.048 | 0.056 |
| 8 | 0.347 | 0.048 | 0.047 | 0.048 | 0.05 | 0.047 | 0.045 | 0.044 | 0.043 | 0.044 | 0.047 | 0.055 |
| 4 | 0.336 | 0.049 | 0.052 | 0.05 | 0.05 | 0.046 | 0.047 | 0.045 | 0.046 | 0.046 | 0.049 | 0.057 |
| 2 | 0.368 | 0.056 | 0.054 | 0.055 | 0.061 | 0.05 | 0.051 | 0.046 | 0.046 | 0.046 | 0.05 | 0.056 |
| 0 | 0.341 | 0.252 | 0.228 | 0.241 | 0.218 | 0.206 | 0.179 | 0.118 | 0.092 | 0.045 | 0.047 | 0.051 |

B

***pgl***

Oxacillin ($\mu$g/ml) →

| Fosfomycin ($\mu$g/ml) | 0 | 0.25 | 0.5 | 1 | 2 | 4 | 8 | 16 | 32 | 64 | 128 | 256 |
|---|---|---|---|---|---|---|---|---|---|---|---|---|
| 128 | 0.07 | 0.053 | 0.049 | 0.048 | 0.049 | 0.047 | 0.05 | 0.047 | 0.049 | 0.051 | 0.057 | 0.059 |
| 64 | 0.156 | 0.05 | 0.049 | 0.048 | 0.05 | 0.047 | 0.048 | 0.045 | 0.047 | 0.05 | 0.052 | 0.063 |
| 32 | 0.241 | 0.05 | 0.052 | 0.051 | 0.051 | 0.05 | 0.054 | 0.053 | 0.05 | 0.049 | 0.05 | 0.064 |
| 16 | 0.308 | 0.063 | 0.065 | 0.056 | 0.059 | 0.055 | 0.063 | 0.063 | 0.06 | 0.05 | 0.05 | 0.062 |
| 8 | 0.32 | 0.091 | 0.094 | 0.097 | 0.092 | 0.089 | 0.1 | 0.095 | 0.086 | 0.049 | 0.05 | 0.069 |
| 4 | 0.307 | 0.182 | 0.185 | 0.196 | 0.196 | 0.193 | 0.2 | 0.187 | 0.15 | 0.072 | 0.052 | 0.073 |
| 2 | 0.319 | 0.273 | 0.261 | 0.261 | 0.27 | 0.269 | 0.257 | 0.24 | 0.21 | 0.126 | 0.054 | 0.061 |
| 0 | 0.304 | 0.304 | 0.291 | 0.283 | 0.254 | 0.244 | 0.252 | 0.238 | 0.244 | 0.19 | 0.05 | 0.059 |

**Fig. S7. Mutation of *pgl* significantly increases resistance to a combination of oxacillin and fosfomycin.** Checkerboard titration assays were conducted using fosfomycin and oxacillin with (A) JE2 and (B) *pgl*, grown for 24 h in Mueller Hinton 2% NaCl broth in 96-well plates. The data shown are the $OD_{600}$ values for each well. The experiments were repeated at least three times and the data from a representative 96-well plate is shown. Green shaded boxes indicated wells in which significant growth was measured.

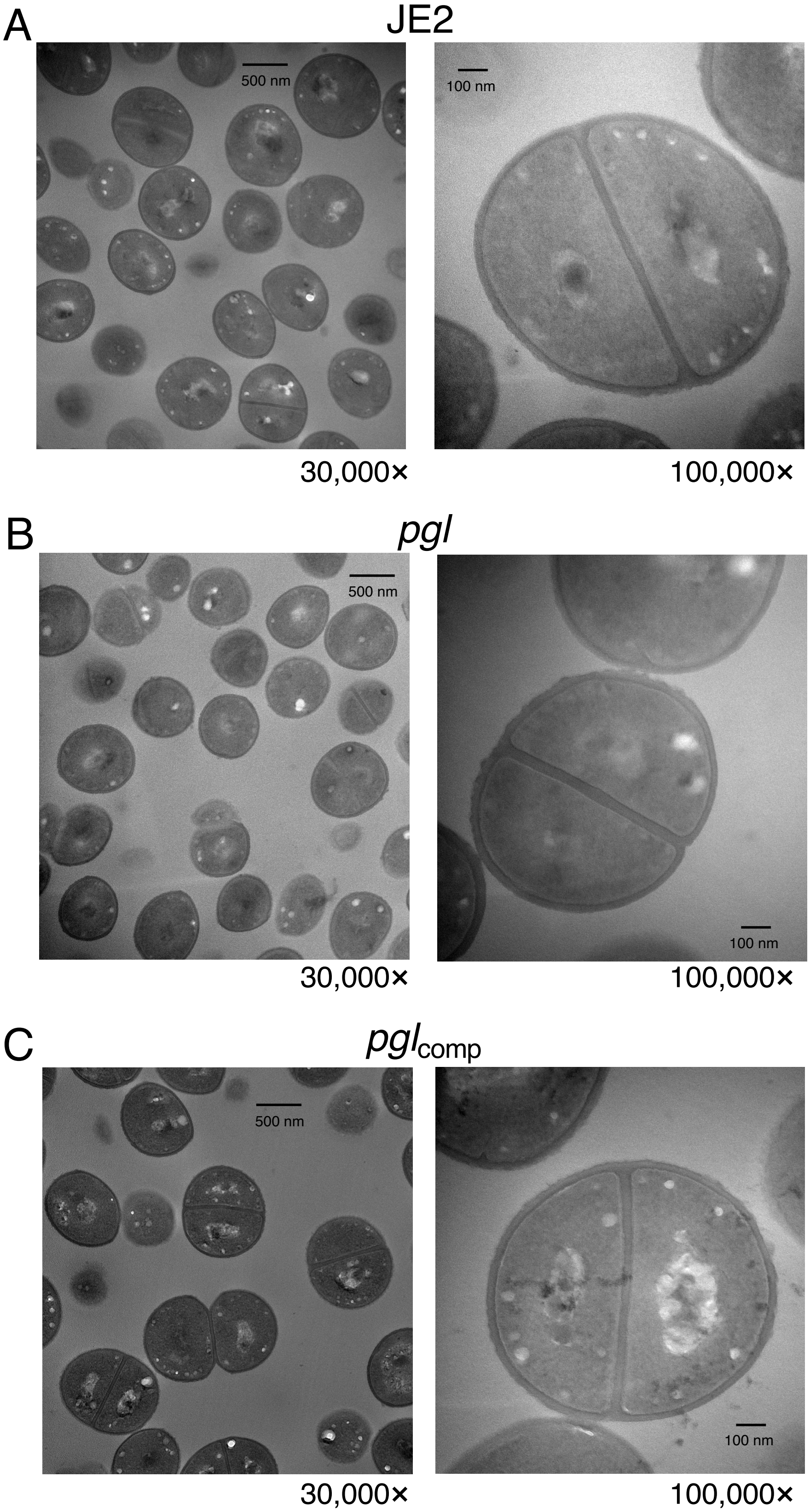


**Fig. S8.** Comparison of wild-type JE2 (A), *pgl* (B) and *pgl*$_{comp}$ (C) cells grown in CDMG without OX using **t**ransmission electron microscopy at 30,000× (left) and 100,000× (right) magnification. Cells were collected from exponential phase cultures grown for 4.5 h in CDMG normalized to $OD_{600}$ = 1 in PBS before being fixed and thin sections prepared. Representative cells from each strain are shown. Scale bars represent 500 nm at 30,000× or 100 nm at 100,000× magnification.

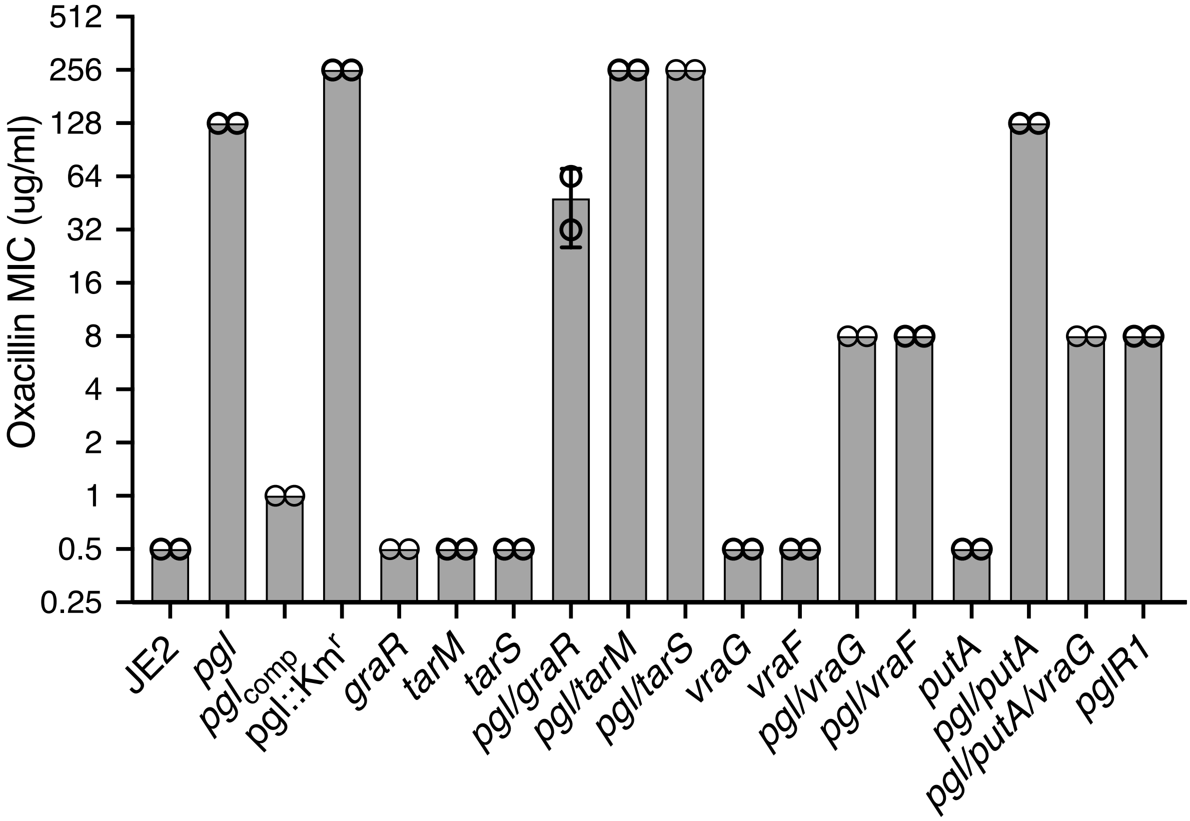


**Fig. S9.** Mutations in *vraF*, *vraG* and *graR* reverse the increased OX minimum inhibitory concentration (MIC) of the *pgl* mutants in CDMG. OX MICs (μg/ml) of JE2, *pgl, pgl*$_{comp}$ *pgl*::Km$^r$, *vraG*, *vraF*, *putA*, *graR*, *tarM*, *tarS*, *pgl*/*tarS*, *pgl*/*tarM*, *pgl*/*vraG*, *pgl*/*vraF*, *pgl*/*putA*, *pgl*/*graR*, *pgl*/*putA*/*vraG* and *pgl*R1 were measured by the broth microdilution method in CDMG. The MIC was measured in two independent experiments for each strain and variation plotted using GraphPad Prism V9 shown.
